# TestNet: A Testing Method for Inferring Microbial Networks with False Discovery Rate Control, for Clustered and Unclustered Samples

**DOI:** 10.1101/2025.04.14.648862

**Authors:** Chang Su, Mengyu He, Vanessa E Van Doren, Colleen F Kelley, Yi-Juan Hu

## Abstract

Most existing methods for inferring microbial networks generate only point estimates of Pearson’s correlations without assessing their significance, and none can account for clustered samples. In this article, we introduce TestNet, a novel method that delivers well calibrated results by controlling the false discovery rate (FDR). We developed a permutation-based procedure to generate valid null replicates that account for compositional effects and extensive zeros in microbiome data, and clustering structures within the samples when present. Our results demonstrate that TestNet is the only method that effectively controls the FDR while maintaining high power across a wide range of scenarios.

## Background

The importance of the human microbiome in human health and diseases has increasingly been recognized. Inference of microbial networks is an important problem in microbiome research, as it reveals interdependencies or interactions among microbial taxa within communities. These interactions include mutualism where taxa help or benefit from each other, competition where they compete for resources, commensalism where one benefits while the others are not affected, and parasitism where one benefits at the expense of another. The constructed microbial networks can subsequently be used to identify keystone taxa, which are highly connected with other taxa and significantly influence microbiome structures and functions. Keystone taxa can be manipulated to treat inflammation or restore community stability [1]. Additionally, microbial networks can be used to infer ecological ‘guilds’, which are groups of taxa with co-abundance behaviors and similar functions. Guilds can serve as units for microbiome data reduction or microbial association analysis with disease outcomes [2]. In this article, we use the term ‘dependence’ broadly to refer to any microbial relationship, including both ‘causality’ and ‘association’.

While absolute abundances of individual taxa are of interest, these values are unobservable in sequencing-based experiments. Techniques such as 16S marker-gene sequencing or shotgun metagenomic sequencing produce taxon count data, where the total counts per sample are considered experimental artifacts. Consequently, these experiments provide information solely about taxon relative abundances, making the data compositional since relative abundances sum to one. Because a change in the relative abundance of one taxon necessarily implies a counterbalancing change in other taxa (i.e., compositional effects), standard correlation analysis of the relative abundances tends to yield negative correlations even if the underlying absolute abundances are independent. The analysis of compositional data typically involves some form of log-ratio transformation of the relative abundances, such as the centered-log-ratio (clr) transformation [3, 4]. Log-ratio transformation effectively removes the compositional effects (as the ratio of relative abundances of two taxa is independent of other taxa), and the transformed relative abundances have been used to make inference about absolute abundances in association studies [5, 6].

In addition, taxon count data exhibit several other complex features. These data are well known to be sparse, with 50–90% zeros. To enable log-ratio transformation, a common practice is to add a pseudo count, typically 0.5 or 1, to the zeros or all entries of the taxa count table [3, 5]. However, the impact of this strategy is often left unevaluated. These data are high-dimensional, with 100–10000 taxa, and it is generally accepted that the dependence structures among the large number of microbial taxa within a community are sparse [7–10]. These data are highly overdispersed due to not only sample heterogeneity but also technical variation in the sequencing workflow, which includes steps such as DNA extraction, PCR amplification, amplicon sequencing, and taxonomy assignment. It is impossible to find a single parametric model that fits every taxon well. These data are also subject to experimental bias because each step in the aforementioned sequencing workflow preferentially measures some taxa over others [11]. Furthermore, microbial taxa within a community have complex interdependencies, and some relationships may be nonlinear [10].

It has become increasingly common to collect repeated (e.g., longitudinal) measures of the microbiome from the same subject [12, 13]. Not only do measures *across* subjects provide information about taxon interdependencies, but measures *within* subjects particularly facilitate this inference, as the large subject heterogeneity as well as all confounding factors between subjects are eliminated in within-subject comparisons.

There are generally two approaches to inferring microbial networks. One approach is based on the marginal dependence between each pair of taxa, independent of all other taxa (e.g., SparCC [7]). The other approach is based on the conditional dependence between each pair of taxa while controlling for all other taxa (e.g., SpiecEasi [14]). While the first approach is unable to differentiate between direct and indirect dependencies, the second one requires more assumptions, such as the normality of data and the relationships among all taxa, and is more computationally complex. Here, we focus on the initial scan of taxon-taxon relationships based on marginal dependence and aim to provide robust inference in the context of complex microbiome data. More precise microbial relationships can be delineated using more complex models after sub-networks are identified and the data dimension is reduced.

Existing methods for inferring marginal dependencies primarily focus on estimating the *basis* correlations, specifically Pearson’s correlations of the underlying (log) absolute abundances. Then, they impose sparsity in dependencies through thresholding or penalization to identify only a few significant dependent pairs. SparCC [7] solves for the basis correlations using an iterative algorithm and then declares dependencies with a threshold of 0.3; the resulting correlations are claimed to be similar to the sample correlations of clr-transformed data when the number of taxa is large. CCLasso [8] estimates the basis covariances (and then the basis correlations) by minimizing the weighted least squares loss between the sample covariance matrix of clr data and the basis covariance matrix, combined with the *L*_1_ penalty to explicitly exploit the sparse dependence assumption. COAT [9] estimates the basis covariances by directly thresholding the sample covariances of clr data, while providing theoretical justification. These methods ignore the influence of sequencing variation (i.e., errors throughout the sequencing workflow) and the impact of their treatments of zeros, which may cause their estimators to be inconsistent. More importantly, none of these methods generate *p*-values or control for any error rate, leading to findings that are not calibrated. In fact, their sensitivity to detecting true correlations may decrease as the sample size increases.

The recently developed method SECOM [10] ‘normalizes’ the observed count data to recover the underlying absolute abundances. These estimates are then used to calculate Pearson’s correlations for measuring linear dependencies and distance correlations [15] for measuring nonlinear dependencies. When their weights are set to be equal, the normalized counts are essentially clr-transformed data. SECOM handles zero counts by restricting its analysis to complete cases when both taxa in the pair have non-zero counts. Unlike other methods, SECOM generates *p*-values, which are based on the *t*-test for Pearson’s correlations and permutation tests for distance correlations. However, there is no theoretical justification for either test. Additionally, while SECOM accounts for experimental bias and sequencing variation, it does not accommodate clustering structures in the samples.

In this article, we address a significant gap in microbial network analysis by introducing a novel testing method called TestNet. We first demonstrate that the basis correlation coefficients are unidentifiable, which motivated us to focus on testing instead of estimation. We propose using the sample Pearson’s covariance and distance covariance of clr data as statistics for testing linear and nonlinear dependencies, respectively. We also propose a strategy for handling zero counts. Furthermore, we develop a permutation-based procedure for generating null replicates that accounts for compositional effects, extensive zero counts, and clustered samples. Finally, we construct an omnibus test that integrates results from both linear and nonlinear tests, enhancing power to detect general forms of dependencies. We evaluate TestNet against existing methods using extensive simulation studies and apply TestNet to two real microbiome studies: one consisting of independent samples and the other consisting of clustered samples.

## Results

### Performance of TestNet and existing methods

Using simulated data, we compared the performance of TestNet with SECOM, SparCC, CCLasso, and COAT. We applied TestNet and SECOM to the simulated data to infer both linear and nonlinear dependencies, and SparCC, CCLasso, and COAT to infer linear dependencies only. For SparCC and CCLasso, we applied a threshold of 0.3 to their estimated correlations. COAT has its own internal thresholding, so no additional thresholding was performed on its estimated correlations. SECOM produced *p*-values, which were used to detect dependencies by the BH procedure. All details regarding the simulation study design are provided in the Methods.

We evaluated the methods using the following metrics: false positive rate (FPR, i.e., type I error) at the nominal level of 0.005 (as used in [10]), empirical FDR at the nominal level of 10%, sensitivity (i.e., recall) at the nominal FDR of 10%, and area under the precision-recall curve (AUC, sometimes referred to as PRAUC in the literature), where precision equals 1 − FDR. While the FPR offers insights into error rate control for individual tests, the FDR is more appropriate for managing errors in multiple testing scenarios. The AUC assesses the overall performance of a method across all nominal FDR levels. All results were based on 100 data replicates, except for the global null case (i.e., Identity covariance structure), where the FDR in each replicate was either 0 or 1 (i.e., exhibiting high variability) and thus the results were based on 1000 replicates to obtain more stable estimates. Note that for SparCC, CCLasso, and COAT, the FPR, FDR and sensitivity were calculated based on their detections, which are independent of the nominal levels set here; their AUC was calculated based on their estimated correlations without thresholding.

We considered a wide range of sample sizes *n*, between 50 and 1000. We extended the upper limit to 1000 to account for the availability of large-scale microbiome studies [20, 21]. We began with cases where the data were simulated with independent samples, *J* = 100 taxa, Normal distributions, and no experimental bias. The results of inferring linear and nonlinear dependencies, based on data with the stringent and lenient filters, are displayed in Figures 1, 2, 3, and S1, respectively. In all these cases, TestNet yielded the most accurate empirical error rates, both FPR and FDR, which closely aligned with their respective nominal levels. Only when *n* became very large (e.g., 1000) did TestNet produce slightly inflated FPR and FDR (e.g., under AR4), as the bias term ℬ_2_ becomes more pronounced. Under the Identity covariance structure, where ℬ_2_ = 0, TestNet exhibited no inflation in FPR regardless of *n*. TestNet showed the highest sensitivity among all methods, with the exception of COAT, which suffered from severely inflated error rates. Additionally, TestNet achieved the highest AUC. The only exception was its linear test under the U-shaped structure, where any linear test is expected to have little power (Figure 1).

**Figure 1.**
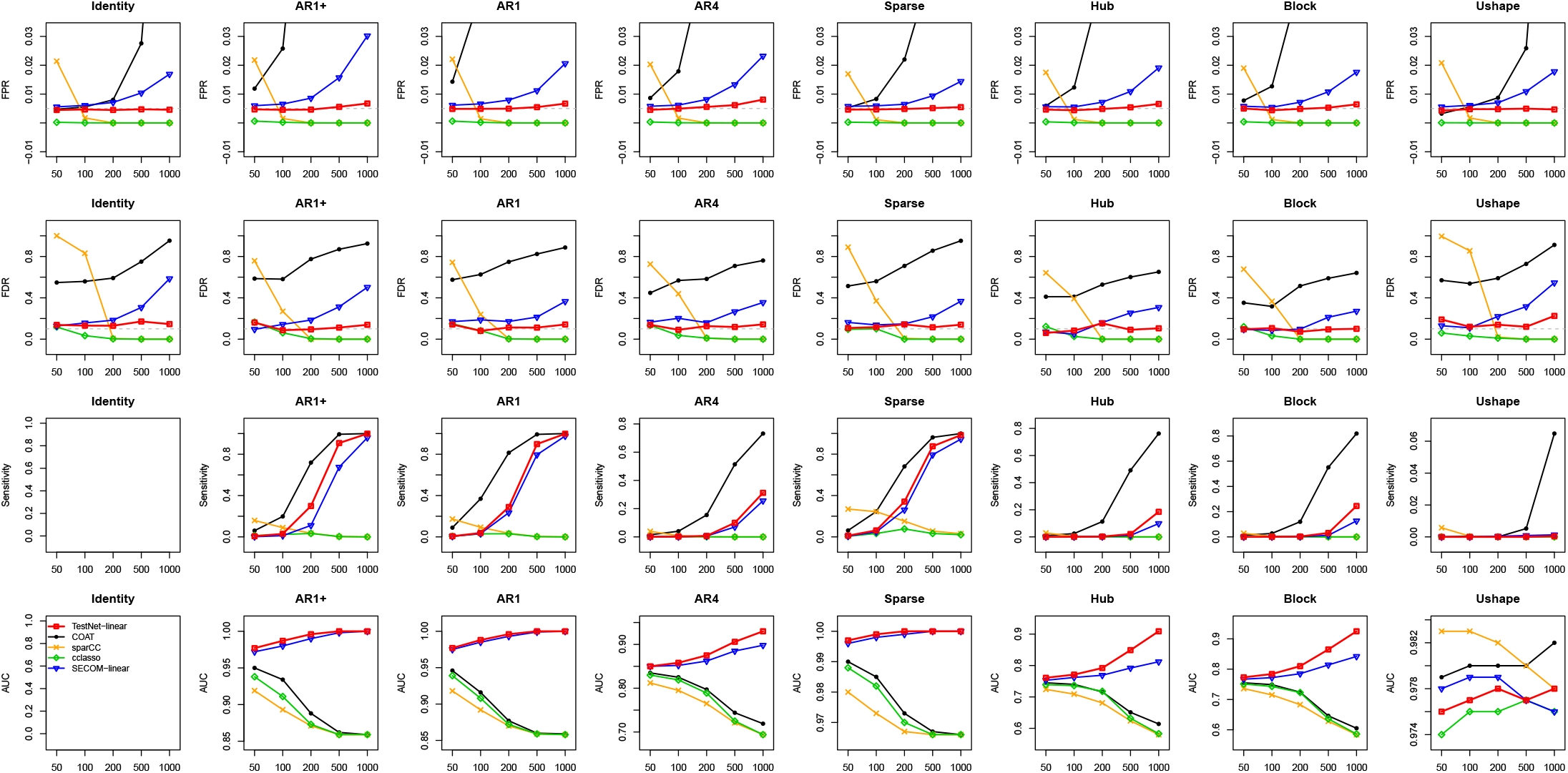
Results on inference of linear dependencies based on data simulated with sample sizes (x-axis) varying from 50 to 1000, independent samples, *J* = 100, Normal distributions, and no experimental bias, and filtered with the stringent criterion. ‘FPR’ is the false positive rate (i.e., type I error) at the nominal level of 0.005 (gray dashed line). ‘FDR’ is the empirical false discovery rate at the nominal level of 10% (gray dashed line). ‘Sensitivity’ is calculated based on the nominal FDR of 10%. ‘AUC’ is the area under the precision-recall curve.

**Figure 2.**
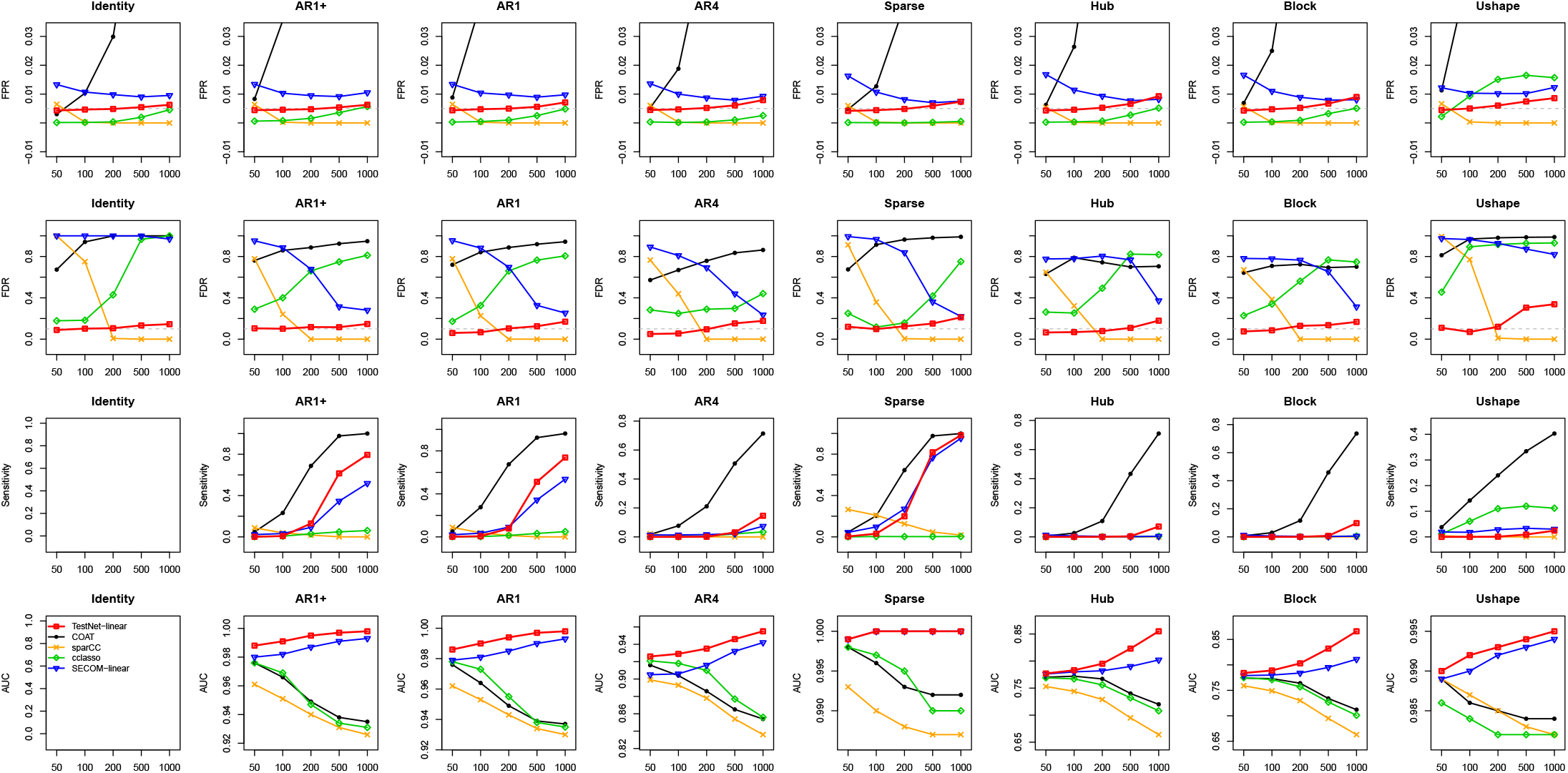
Results on inference of linear dependencies based on data filtered with the lenient criterion. Refer to the caption of Figure 1 for additional information.

**Figure 3.**
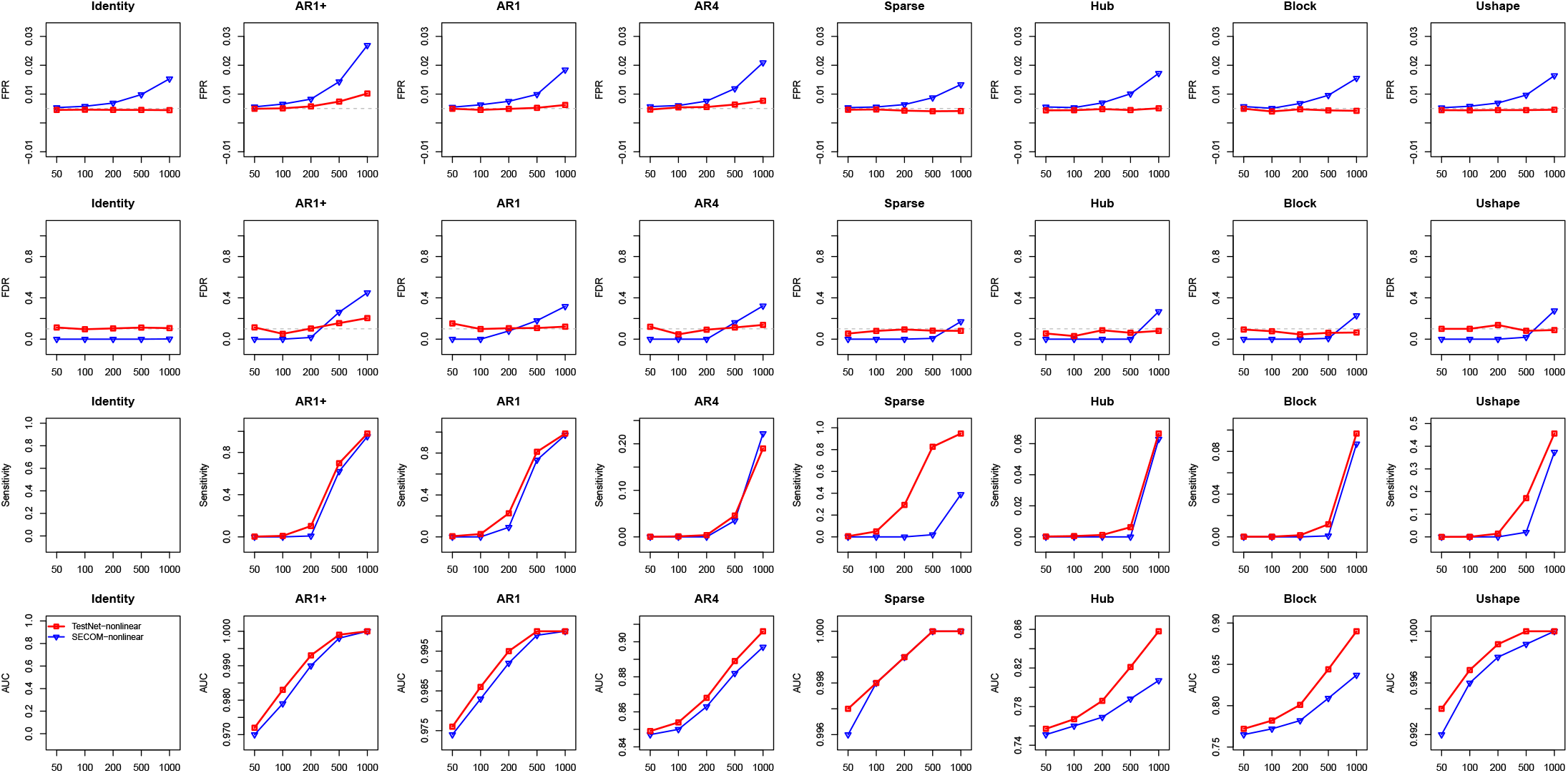
Results on inference of nonlinear dependencies based on data filtered with the stringent criterion. Refer to the caption of Figure 1 for additional information.

Among the existing methods, SECOM demonstrated performance most similar to TestNet, though it had lower sensitivity and AUC. However, SECOM failed to control error rates in many instances, especially when *n* was large or when testing linear dependencies at rare taxa due to the use of a lenient filter. SparCC exhibited low sensitivity and the lowest AUC, with high error rates when *n* was small. Its sensitivity even decreased as *n* increased, as its estimated correlations converged to values lower than the true correlations. CCLasso generally yielded low error rates and sensitivity, although its FDR was sometimes high (e.g., in Figure 2). Conversely, COAT typically had high error rates and sensitivity, but its AUC was low.

These conclusions generally hold when log *Z* were sampled from Gamma distributions (Figures S2 and S3) or when the number of taxa *J* was set to 50 (Figures S4 and S5). As expected, TestNet’s inflation of error rates under large samples sizes slightly worsened when *J* became unrealistically small.

### Adding experimental bias

The results for simulated data with experimental bias are displayed in Figures S6 and S7. Compared to Figures 1 and 2, experimental bias had minimal effects on the results of TestNet and other existing methods. This outcome is expected, as all methods are based on or closely related to Pearson’s covariance or distance covariance of clr-transformed data, both of which have been shown earlier to be invariant to multiplicative experimental bias.

### Clustered data

With a fixed cluster size of four, we varied the number of clusters between 13 and 250 to achieve sample sizes between 52 and 1000. We applied TestNet with three different permutation schemes: shuffling data both between and within clusters (referred to as TestNet-b-w), shuffling data between clusters only (TestNet-b), and shuffling all samples irrespective of the clustering structure (TestNet-free). The results for data filtered with the stringent and lenient criteria are shown in Figures 4 and S8, respectively. As expected, TestNet-b-w demonstrated the best performance among the three versions of TestNet. TestNet-free yielded inflated error rates, with a level of inflation similar to that of SECOM (when *n* was small), since SECOM also employed the free permutation scheme regardless of the clustering structure. TestNet-b, without the within-cluster shuffling, produced lower sensitivity than TestNet-b-w, particularly under the Sparse structure.

**Figure 4.**
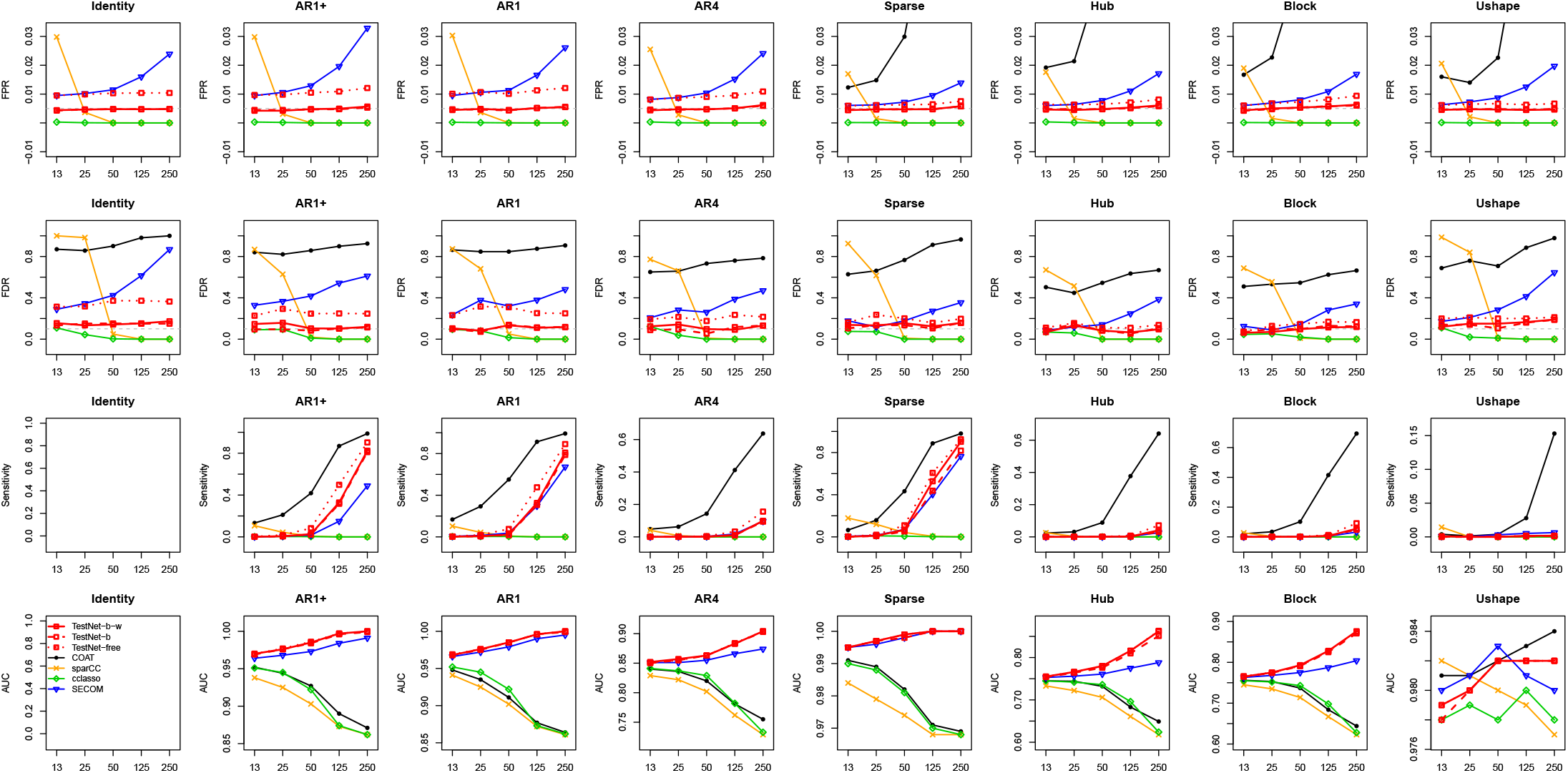
Results on inference of linear dependencies based on clustered data simulated with the number of clusters (x-axis) varying from 13 to 250 and four samples in each cluster, and filtered with the stringent criterion. Refer to the caption of Figure 1 for additional information.

### Omnibus test

The results of the TestNet omnibus test, along with its linear and nonlinear test results, for data filtered with the stringent and lenient criteria are presented in Figures S9 and S10, respectively. The omnibus test controlled both error rates, FPR and FDR, and consistently tracked either the linear or nonlinear test, whichever demonstrated superior performance in sensitivity and AUC.

### Longitudinal rectal mucosal microbiome data

The first real dataset we analyzed was generated from a longitudinal clinical study conducted by our research team [22] to investigate microbiome responses to experimentally induced rectal mucosal injury in men who have sex with men (MSM). At the baseline visit, 19 MSM underwent a rectal mucosal injury using biopsy forceps approximately 6 cm from the anal verge. Participants returned for follow-up visits on days 2, 5, and 8. Rectal mucosal secretions were collected at all four visits for microbiome composition profiling. These samples were sequenced by Illumina MiSeq for the 16S rRNA gene, which data were then summarized into a taxa count table at the genus level. Our goal was to identify inter-genus dependencies within the rectal mucosal microbiome community of these MSM.

There were 372 genera with at least one read across the 76 samples. The library sizes were adequate, ranging from 12,601 to 353,656, with a median of 133,932 and a mean of 135,771. We applied the stringent filter to exclude genera with average relative abundances less than 0.1% across the 76 samples, resulting in 106 genera for network analysis. The ordination plot in Figure 5 shows that the four samples from each subject tend to cluster together, indicating that microbiome profiles are more similar within subjects than across subjects. Therefore, it is important to account for this clustering structure when inferring the microbial network.

**Figure 5.**
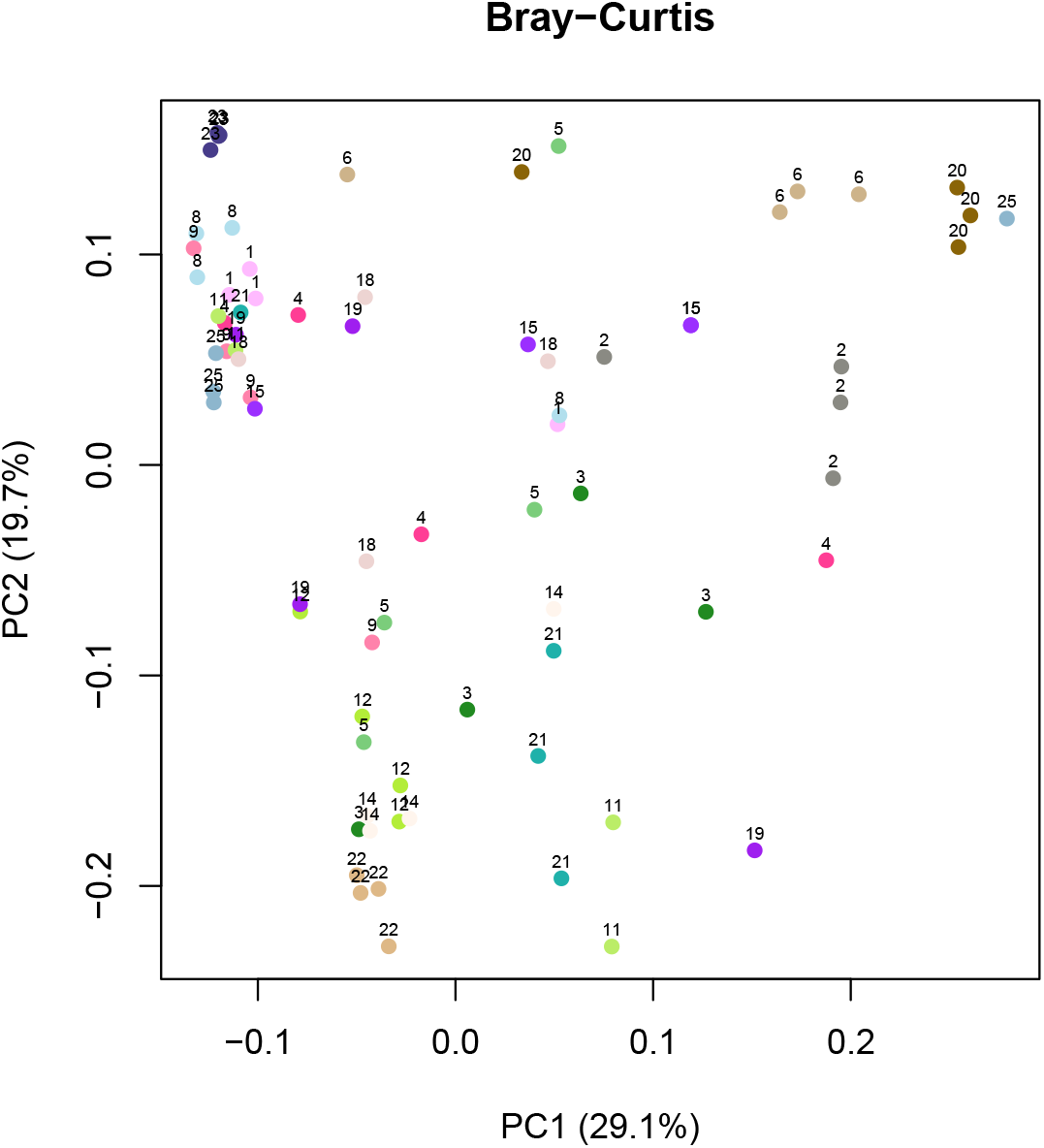
Ordination of the longitudinal gut microbiome samples based on the Bray-Curtis distance. Samples from the same subject are marked with the same color and the same subject ID.

We applied TestNet and other methods in the same manner as in the simulation studies. Specifically, we used TestNet with both between- and within-cluster permutations to preserve the clustering structure. The other methods were applied without considering the clustering structure. The numbers of dependent pairs of genera detected by these methods are displayed in Table 1. TestNet, using the omnibus test, detected 240 general dependencies at a nominal FDR level of 0.5%. With this nominal FDR level, we expect the detected dependencies to include approximately one false positive. As expected, the linear and nonlinear tests of TestNet detected less dependencies, 213 and 223 respectively, than the omnibus test (240). However, the three numbers of detections do not differ significantly, as the linear and nonlinear tests are inherently overlapping: their test statistics can capture both types of relationships. SECOM applied its linear and nonlinear tests separately to each pair of taxa. The linear test detected 210 dependencies, while the nonlinear test detected none–a discrepancy that is difficult to reconcile given the overlap of the two test statistics. In addition, applying the linear and nonlinear tests separately made it challenging to control the overall error rate. The other methods detected linear dependencies only. The number of detections varied dramatically between methods, from 5,010 to 768, highlighting the importance of selecting the appropriate method. In particular, some of these numbers, such as 1,146 and even 5,010, are unreasonably high, as there are only 5,565 tests in total. Note that none of the existing methods can control the FDR, especially in the presence of clustered samples.

**Table 1:**
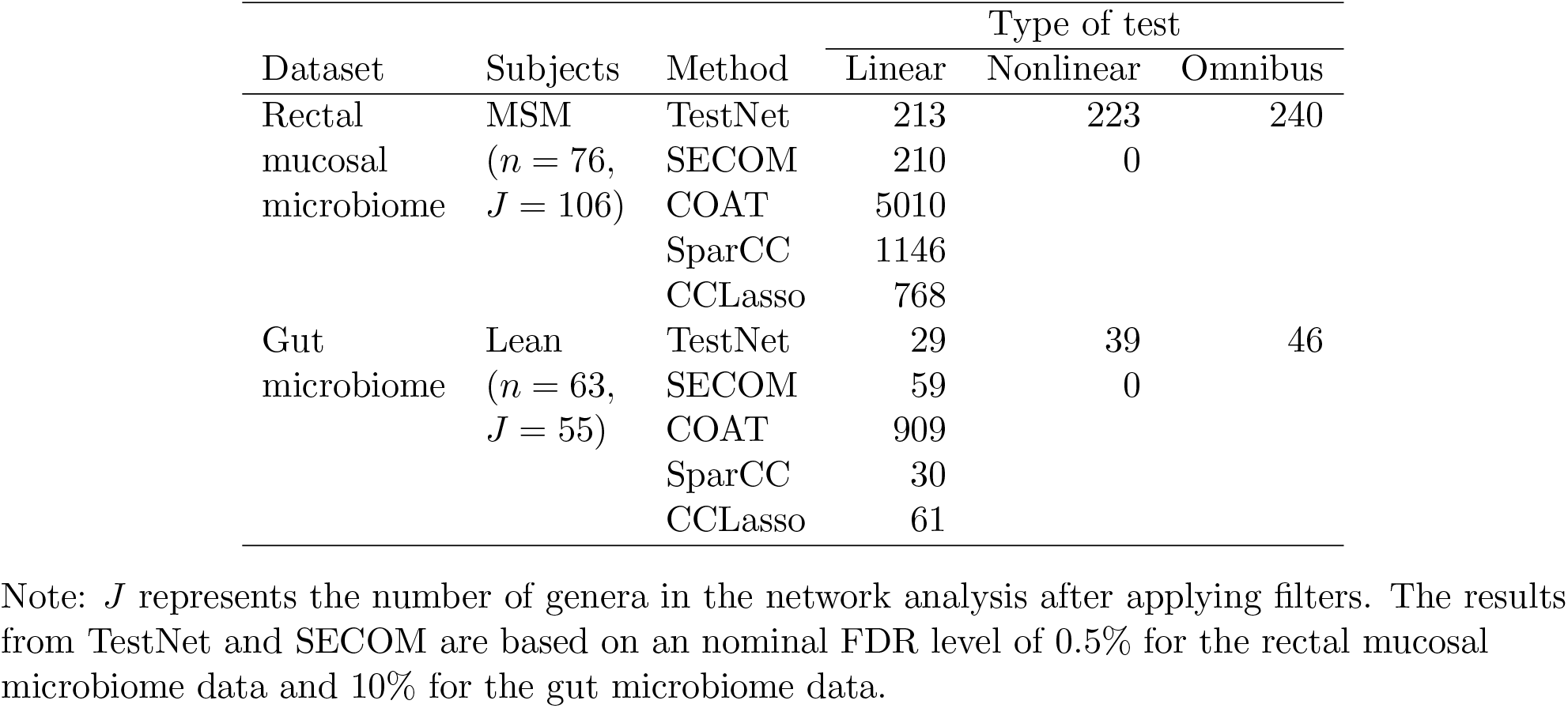
Number of dependent pairs of taxa detected in the real datasets.

| Dataset | Subjects | Method | Type of test |  |  |
| --- | --- | --- | --- | --- | --- |
|  |  |  | Linear | Nonlinear | Omnibus |
| Rectal mucosal microbiome | MSM<br>($n = 76$,<br>$J = 106$) | TestNet | 213 | 223 | 240 |
|  |  | SECOM | 210 | 0 |  |
|  |  | COAT | 5010 |  |  |
|  |  | SparCC | 1146 |  |  |
|  |  | CCLasso | 768 |  |  |
| Gut microbiome | Lean<br>($n = 63$,<br>$J = 55$) | TestNet | 29 | 39 | 46 |
|  |  | SECOM | 59 | 0 |  |
|  |  | COAT | 909 |  |  |
|  |  | SparCC | 30 |  |  |
|  |  | CCLasso | 61 |  |  |

Note: $J$ represents the number of genera in the network analysis after applying filters. The results from TestNet and SECOM are based on an nominal FDR level of 0.5% for the rectal mucosal microbiome data and 10% for the gut microbiome data.

Thus far, we have detected general dependencies using the TestNet omnibus test. To get further insights into whether each general dependence is roughly linear or nonlinear, we employed a heuristic strategy by comparing the linear and nonlinear test *p*-values. A total of 153 dependencies had linear *p*-values lower than or equal to their nonlinear counterparts, and we classified them as linear dependencies. The remaining 87 dependencies were classified as nonlinear dependencies.

We further visualized the detected (general) dependencies by TestNet as network edges in Figure 6. The most connected genera include *Peptoniphilus* (21 edges), *Prevotella* (19 edges), *Eubacterium hallii group* (19 edges), *Anaerococcus* (17 edges), and *Peptostreptococcus* (15 edges). These genera are all abundant and have previously been associated with receptive anal intercourse (RAI) in [23], suggesting that they are likely keystone taxa exerting a profound influence on the structure of the rectal mucosal microbiome community in MSM. In many subnetworks, their members tend to exhibit (linear) correlations in the same direction, either positive or negative, suggesting that they form ecological guilds that share co-abundance behaviors and similar functions.

**Figure 6.**
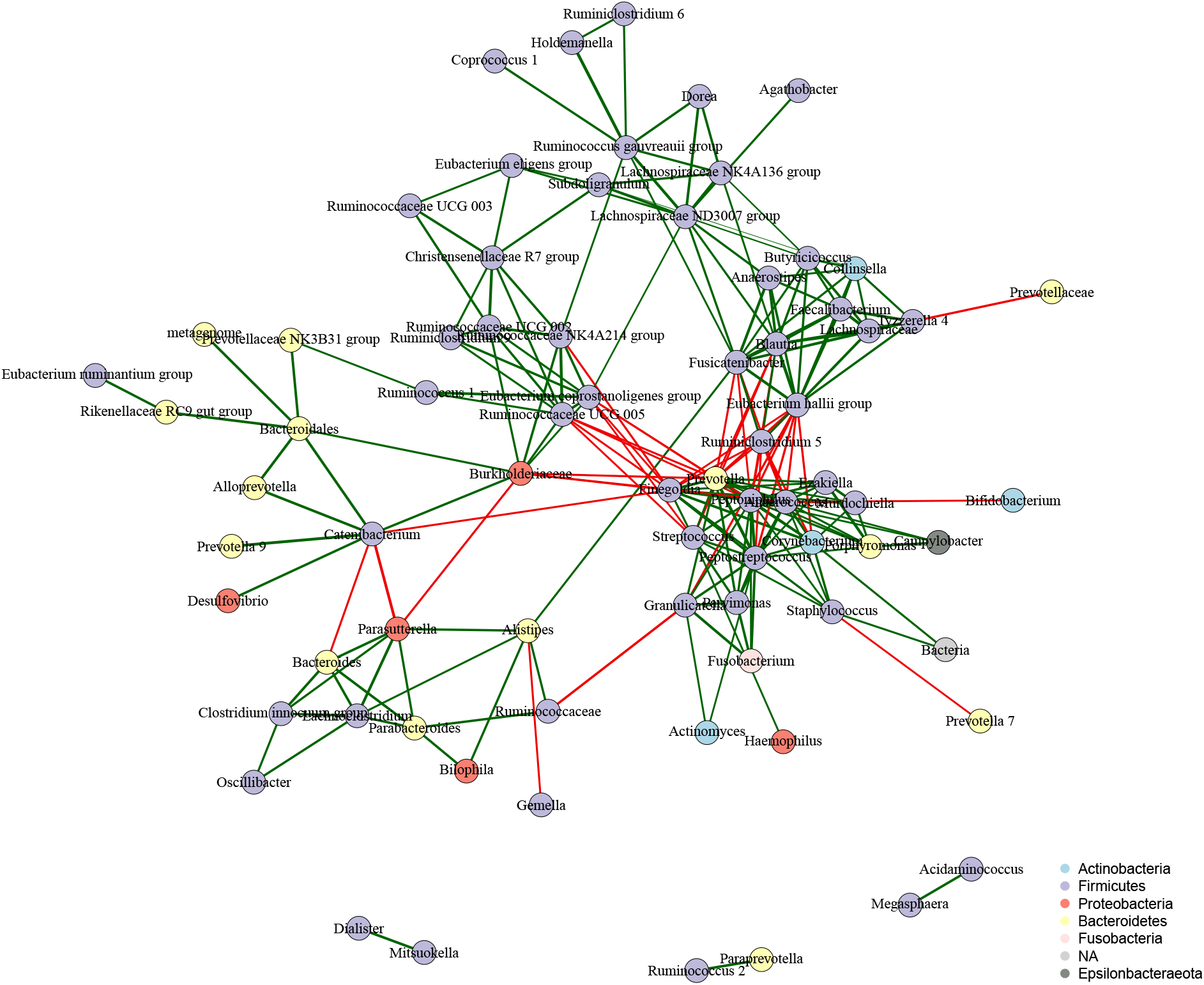
Microbial networks of general dependencies identified by TestNet (the omnibus test) based on the rectal mucosal microbiome data of MSM. Positive and negative linear correlation estimates are displayed as green and red edges, respectively, with the thickness of the edges indicating the magnitude of the linear correlation estimates. Here, the linear correlation estimates were used for illustration purposes only.

### Cross-sectional gut microbiome data

The second real dataset we analyzed was derived from a cross-sectional study [24] that sequenced the stool samples of 63 lean (BMI *<* 25) individuals using 454/Roche pyro-sequencing for the 16S rRNA gene. The sequence data consisted of an average of 9,265 reads per sample (with a standard deviation of 3,864) across 87 genera detected in at least one sample. We applied the lenient filter to focus on the *J* = 55 genera that appeared in at least five samples. Again, our goal was to identify inter-genus dependencies within the gut microbiome community of these lean subjects.

Since the samples were independent, we applied TestNet with free shuffling across all samples. The numbers of detected pairs of taxa are summarized in Table 1. TestNet (omnibus) detected 46 general dependencies (at an FDR of 10%, implying 4–5 false positives), which were classified into 23 linear dependencies and 23 nonlinear ones. Although the other methods were designed for independent samples, their numbers of detections still varied dramatically from method to method. Our simple application of COAT detected 909 correlations. In contrast, the network analysis in the COAT paper [9] retained 254 correlations that were reproduced in at least 85% bootstrap replicates. Despite this, these results may still contain many false positives due to the high error rates of COAT.

The microbial networks inferred by TestNet (omnibus) is visualized in Figure 7. *Bacteroides* is the 2nd most connected taxon in the network. Within this genus, several species, such as *Bacteroides fragilis* and *Bacteroides thetaiotaomicron*, are well established as keystone taxa in the human gut microbiome [1]. The network includes a strong negative correlation between *Bacteroides* and *Prevotella*. The ratio of *Bacteroides* and *Prevotella* has long been used to define gut microbial enterotypes that reflect long-term dietary patterns [24].

**Figure 7.**
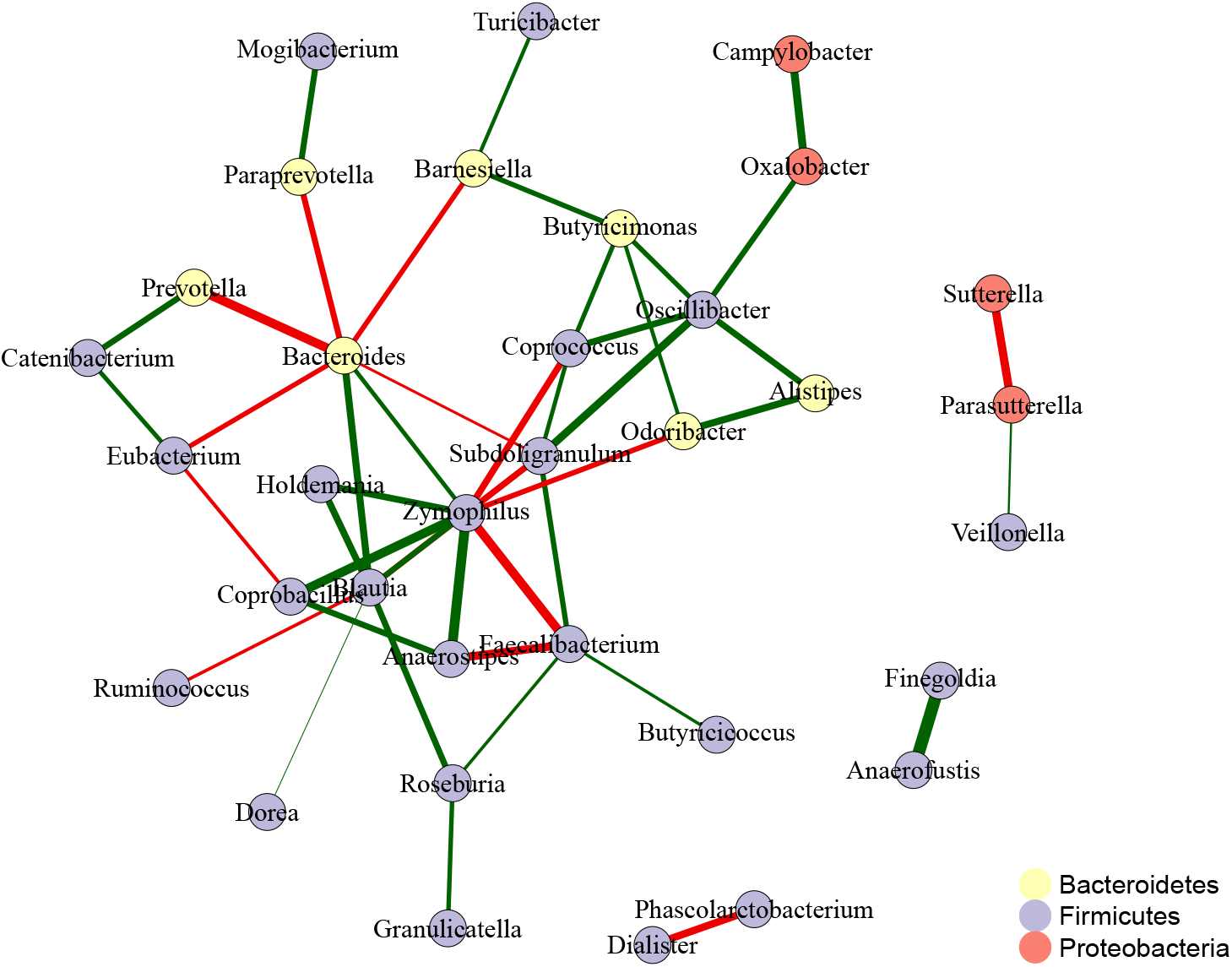
Microbial networks of general dependencies identified by TestNet (the omnibus test) based on the gut microbiome data of lean individuals. Refer to the caption of Figure 6 for additional information.

## Discussion

TestNet shares notable similarities with SECOM. TestNet’s use of the clr transformation aligns with SECOM’s normalization of observed count data. Both methods also measure dependencies using similar metrics: SECOM employs Pearson and distance correlations for linear and nonlinear dependencies, while TestNet uses Pearson and distance covariances. However, there are key differences between the two. SECOM incorporates Pearson and distance variances that are inflated by sequencing variation, which TestNet does not use. While SECOM addresses zero counts by restricting its analysis to complete cases, TestNet leverages the information in zeros by adding a pseudo-count of 0.5 for clr-transformation, followed by mean imputation. Additionally, SECOM overlooks the constraints in the ‘normalized’ data during its *t*-tests and permutation tests, whereas TestNet explicitly accounts for the constraints in the clr data within its permutation procedure. Strikingly, the FDR of SECOM exceeded 80% when constructing a network with 50–100 independent samples that included many rare taxa (e.g., Figure 2), mirroring the scenario observed in the gut microbiome data we analyzed.

TestNet is applicable to compositional data beyond microbiome data, such as gene expression data, brain cell subtypes, and immune cell subsets. It is critical to verify the sparse dependence assumption for each application. Notably, the likelihood of violating this assumption increases as the number of features decreases. To investigate this, we conducted additional simulation studies with the number of features set to *J* = 20 or *J* = 10, and found that the FDR was generally reasonable when sample size was less than 200 (Figures S11–S14).

We have demonstrated several biases in Pearson’s covariance and variance of the clr data, and have provided insights into the cause of each bias. The biases ℬ_1_ through ℬ_4_ all contribute to the inconsistency of correlation estimates. However, only ℬ_2_ may cause FDR inflation for TestNet. Therefore, even when ℬ_1_, ℬ_3_, and ℬ_4_ are significant, such as with a small *J*, TestNet can still control the FDR (e.g., Figures S11–S14 under the Identify structure).

We have implemented our method in the R package TestNet, which is available on GitHub at https://github.com/yijuanhu/TestNet in formats appropriate for Macintosh or Windows. Despite the permutation procedure, TestNet is computationally efficient for data with small sample sizes and very sparse dependencies. For example, using one core of a MacBook Pro laptop (Apple M3 Pro, 18GB memory), it took 27 seconds to analyze a simulated dataset with 100 samples, 100 taxa, and the AR1 structure, 15 seconds to analyze the gut microbiome data, and 10 minutes to analyze the rectal mucosal microbiome data. The extended time for the last analysis is due to the large number of dependencies, requiring over 60,000 permutation replicates. TestNet could be improved by splitting the sequential permutation procedure into blocks of a fixed number of permutations and parallelizing the permutations within each block.

## Conclusions

We have introduced TestNet, the first testing method for inferring microbial networks that provides calibrated results by controlling the FDR. TestNet is highly robust and performs well even with a large number of zeros in the data. TestNet is also highly versatile: it accommodates both independent and clustered samples, offers separate linear and nonlinear tests, and includes an omnibus test that eliminates the need to pre-specify the type of relationship.

## Methods

We consider a microbial community with *J* taxa. Let *Z*_*j*_ be the underlying absolute abundance of taxon *j*, and 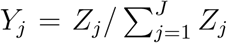 be the underlying relative abundance, both of which are unobserved. Let *X*_*j*_ be the observed read count, whose distribution is governed by *Y*_*j*_ and library size *l* (i.e., sequencing depth). We calculate the observed relative abundance as 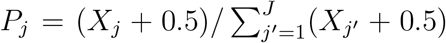, by adding a pseudo count of 0.5 to all counts. For a general variable *W*_*j*_, which can be *Z*_*j*_, *Y*_*j*_ or *P*_*j*_, the clr transformation of *W*_*j*_ is denoted by 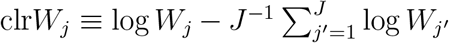. Our method is based on two key facts: (1) clr*Z*_*j*_ = clr*Y*_*j*_, and (2) *P*_*j*_ is a consistent estimator for *Y*_*j*_ (in the absence of experimental bias). Additionally, our method relies on the following two assumptions:

- Sparse Dependence Assumption:

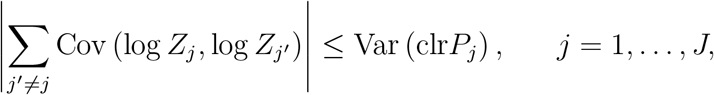

where all Var (clr*P*_*j*_)’s are bounded.
- Independence Assumption: *X*_*j*_s across taxa are independent given (*Y*_1_, *Y*_2_, …, *Y*_*J*_).

The Sparse Dependence Assumption is more easily satisfied in the presence of large overdispersion arising from sequencing variations. In this case, large overdispersion becomes a bless rather than a curse. The Independence Assumption ensures that sequencing variations across taxa are independent. In the subsequent text, we omit the index *i* for samples in general, except for a few scenarios.

### Test statistic for linear dependencies

As in all the aforementioned existing methods, the linear dependence of interest is Pearson’s covariance Cov(log *Z*_*j*_, log *Z*_*k*_) for *j* ≠ *k*. It is related to Cov(clr*Z*_*j*_, clr*Z*_*k*_) through the equation

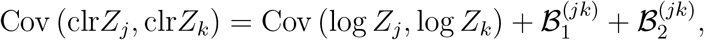

where

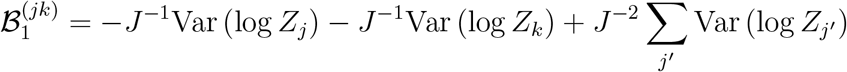

and

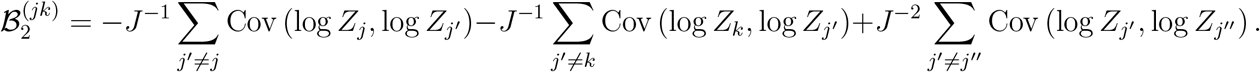

We omit the superscript (*jk*) in 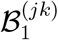 and 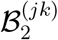 in general unless otherwise specified. We refer to ℬ_1_ and ℬ_2_ as bias terms, which arise from using relative abundances *Y*_*j*_ instead of absolute abundances *Z*_*j*_, since clr*Z*_*j*_ = clr*Y*_*j*_. In particular, ℬ_2_ exclusively accounts for the portion attributable to taxon-taxon dependencies. Since Var(clr*P*_*j*_) is bounded, Var(log *Z*_*j*_) is also bounded due to their relationship in Equation (2) below. Consequently, ℬ_1_ = *O*(*J*^−1^) is negligible as long as *J* is sufficiently large. Similarly, under the Sparse Dependence Assumption, ℬ_2_ = *O*(*J*^−1^) is also negligible. Next, we substitute the observed relative abundances *P*_*j*_ for *Y*_*i*_ in Cov(clr*Y*_*j*_, clr*Y*_*k*_) to obtain Cov(clr*P*_*j*_, clr*P*_*k*_) and calculate the sample-based statistic, 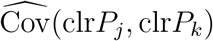, as our test statistic. In fact, all existing methods are closely related to this statistic.

The replacement of *Y*_*i*_ with *P*_*i*_ introduces another source of bias that is often overlooked by existing methods. To demonstrate this, we use conditional expectation to establish the relationship between Cov(clr*P*_*j*_, clr*P*_*k*_) and Cov(clr*Y*_*j*_, clr*Y*_*k*_) for *j*≠ *k*, conditioning on Y = (*Y*_1_, *Y*_2_, …, *Y*_*J*_):

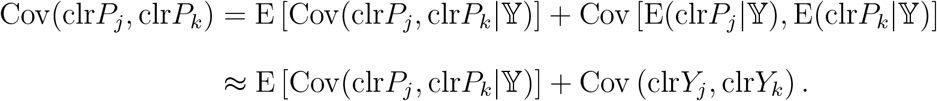

The last step follows from the delta method, which provides an approximation accurate to the order of *O*(*l*^−1^). Recall that *l* denotes the library size, which is at least in the range of thousands. The term ℬ_3_ ≡ E [Cov(clr*P*_*j*_, clr*P*_*k*_| 𝕐)] represents the bias caused by sequencing variation (when the underlying relative abundances are fixed). We derived in Supplementary Text A that, under the Independence Assumption,

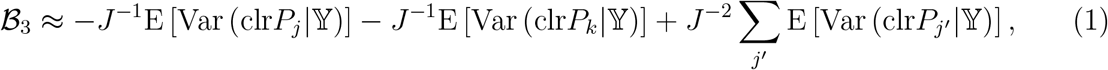

which indicates that B_3_ is an *O*(*J*^−1^) term.

In summary, the overall bias in Cov(clr*P*_*j*_, clr*P*_*k*_) relative to Cov(log *Z*_*j*_, log *Z*_*k*_) is ℬ_1_ +ℬ_2_ + ℬ_3_. Thus, 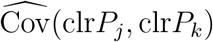 is not a consistent estimator for Cov(log *Z*_*j*_, log *Z*_*k*_) as the sample size *n* goes to infinity. Nevertheless, 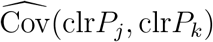 makes a reasonable statistic for testing linear dependencies, since the overall bias is an *O*(*J*^−1^) small term.

### Correlation coefficients cannot be identifiable due to sequencing variation

Although sequencing variation has only a small impact on the covariance, it tends to have a much larger effect on the *variance* and hence the *correlation*. We use the conditional expectation again to establish the relationship between Var(clr*P*_*j*_) and Var(clr*Y*_*j*_):

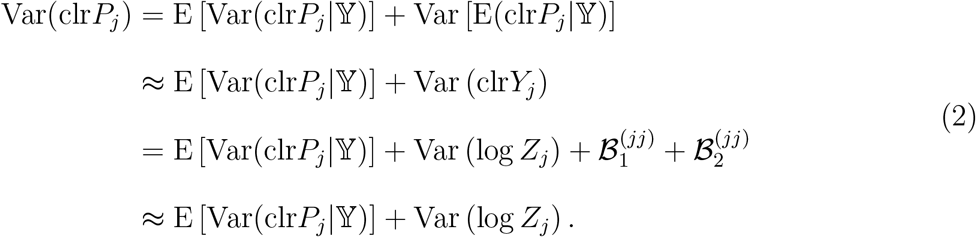

The last approximation follows from the fact that both 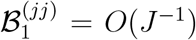 and 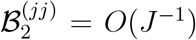 are negligible. If the variance in read count *X*_*j*_ due to sequencing, Var(*X*_*j*_| 𝕐), is at least *O*(*l*^2^) (i.e., it does not approach zero when divided by *l*^2^ as *l* approaches infinity), the bias term B_4_ ≡ E [Var(clr*P*_*j*_| 𝕐)] = E [Var{clr(*X*_*j*_ + 0.5)| 𝕐}] remains significant regardless of the value of *l*. More details can be found in Supplementary Text B. This condition holds for read count data with some level of overdispersion, such as the Negative-Binomial data, though it represents a very moderate level of overdispersion. This bias causes the variance Var(clr*P*_*j*_) to substantially overestimate Var(log *Z*_*j*_) and, consequently, the correlation Cor(clr*P*_*j*_, clr*P*_*k*_) to markedly underestimate Cor(log *Z*_*j*_, log *Z*_*k*_). This result has important implications for methods that rely on thresholding 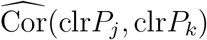, as they tend to miss true correlations. Since the correlation coefficient is unidentifiable, we use 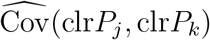 as a test statistic and focus on testing its deviation from the null hypothesis.

### Test statistic for nonlinear dependencies

Following SECOM, we adopt *distance covariance* [15] to measure nonlinear dependencies and denote the distance covariance of two variables, *W*_1_ and *W*_2_, by dCov(*W*_1_, *W*_2_). By definition, distance covariance measures the difference in the joint characteristic function of *W*_1_ and *W*_2_ with and without the assumption of independence between *W*_1_ and *W*_2_. The sample distance covariance, denoted by 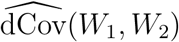, is calculated as the sample Pearson’s covariance between the (centered) pairwise distances among the *W*_1_ data and the distances among the *W*_2_ data. Specifically, denote the *W*_1_ data on *n* samples by {*W*_1*i*_, *i* = 1, …, *n*}, and compute the *n* × *n* Euclidean distance matrix {*d*_1,*ii*_*′*} = {|*W*_1*i*_ − *W*_1*i*_*′* |} and then the centered distance matrix 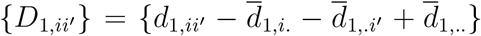, where 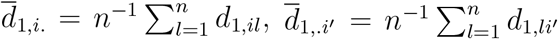, and 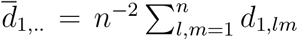. Similarly, compute the distance matrix and centered distance matrix based on the *W*_2_ data as {*d*_2,*ii*_*′*} and {*D*_2,*ii*_*′*}. Finally, calculate 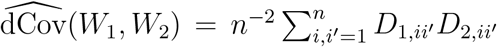. Unlike Pearson’s covariance, distance covariance is always non-negative. A zero distance covariance indicates independence of two random variables, whereas a zero Pearson’s covariance does not does not necessarily imply independence unless the variables are Gaussian-distributed. Although distance covariance can capture more general patterns of dependencies including linear ones, it is less efficient than Pearson’s for capturing linear dependencies. More details about distance covariance can be found in [15, 16]. Our distance covariance of interest is dCov(log *Z*_*j*_, log *Z*_*k*_). Following the procedure in the linear case, we replace log *Z*_*j*_ with clr*Z*_*j*_, equate clr*Z*_*j*_ with clr*Y*_*j*_, and substitute clr*P*_*j*_ for clr*Y*_*j*_. We obtain the test statistic for nonlinear dependencies as 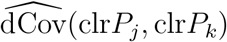. Like its linear counterpart, distance covariance and variance are subject to bias due to data compositionality and sequencing variation. However, it remains unclear how to derive these bias terms in closed forms, so we resort to simulation studies for their evaluation.

### Handling zeros

The use of pseudo-counts (e.g., 0.5) to address zeros, while common in microbiome data analysis, introduces a problem that, fortunately, has a straightforward solution. For a zero entry of the taxa count table corresponding to sample *i* and taxon *j*, i.e., *X*_*ij*_ = 0, the clr-transformed value is clr*P*_*ij*_ = log 0.5 − *J*^−1^ ∑_*j*_*′* log(*X*_*ij*_*′* + 0.5). This value is invariant across taxa but varies across samples, which tends to create spurious associations between pairs of taxa, especially when both taxa in a pair have extensive zeros. To eliminate these spurious associations, we propose replacing such clr*P*_*ij*_ values with their taxon-specific means before calculating the test statistics, where the taxon-specific means are based on the clr*P*_*ij*_ values at zero entries only. This modification effectively breaks the spurious associations at zero entries while maximally preserving the orders of clr*P*_*ij*_ values between non-zero and zero entries. We denote the modified clr*P*_*ij*_ value by mclr*P*_*ij*_ and redefine the test statistics as 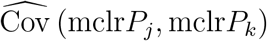 and 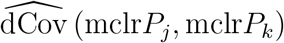.

### Impact of experimental bias

In the presence of experimental bias, the underlying relative abundance that we aim to ‘measure’, denoted by 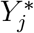, differs from the true relative abundance *Y*_*j*_. We assume that 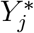 is related to *Y*_*j*_ and the taxon-specific bias factor *γ*_*j*_ through the WMC model [11]:

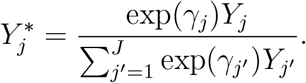

It follows that their clr-transformed values differ only by a taxon-specific constant:

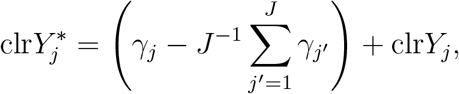

which leads to 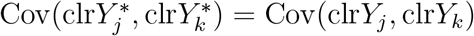 and 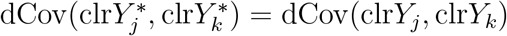. Since *P*_*j*_ is now a consistent estimator for 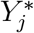 (not *Y*_*j*_), our test statistics still target at the desired dependencies of interest.

### Permutation-based inference

--

Asymptotic inference using 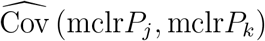 and 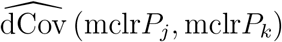 as test statistics may not work well for several reasons. Firstly, most microbiome studies have small sample sizes ranging from 50 to 200. Secondly, the distribution of mclr*P*_*j*_ values still tends to be skewed, although less skewed than the original relative abundances *P*_*j*_. Thirdly, the mclr*P*_*j*_ distribution has a point mass resulting from the modification at zeros. Therefore, we rely on permutation for inference. The permutation procedure is based on shuffling the clr*P*_*j*_ values for each taxon, as they have been adjusted to eliminate compositional effects and are therefore independent across taxa under the null hypothesis. The rationale for not shuffling the mclr*P*_*j*_ values will be explained later.

Recall that Cov(clr*P*_*j*_, clr*P*_*k*_) has a non-zero value ℬ_1_ + ℬ_2_ + ℬ_3_ under the null hypothesis that Cov(log *Z*_*j*_, log *Z*_*k*_) = 0. Therefore, the covariance calculated from the permutation replicates should have the same non-zero mean to ensure their validity. Let 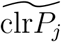 be the shuffled

clr*P*_*j*_ across samples, where shuffling is done independently for different taxa to generate a permutation replicate. Unfortunately, the naive covariance 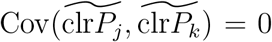 due to the independence of 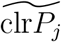 and 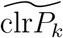 for any *j* ≠ *k*. Note that 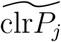 across taxa no longer sums to zero as the original clr*P*_*j*_ does, so we re-center 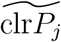 to obtain a modified covariance

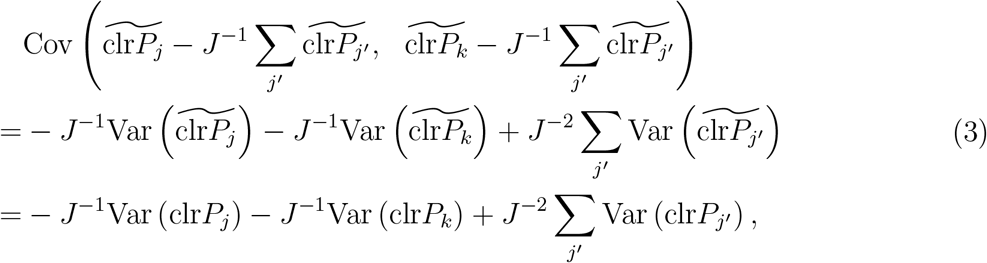

where the last equation follows from the fact that shuffling preserves variance. Using (2), we can express (3) as ℬ_1_+ℬ_3_. This value nearly matches Cov(clr*P*_*j*_, clr*P*_*k*_) = ℬ_1_+ℬ_2_+ℬ_3_ under the null hypothesis, except for the loss of the B_2_ term. Recall that B_2_ = 0 under the Identity covariance structure and ℬ_2_ = *O*(*J*^−1^) in general cases satisfying the Sparse Dependence Assumption. Meanwhile, the sample covariance 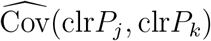, which is closely associated with the final test statistic, has a standard error of *O*(*n*^−0.5^). When *n* is small relative to *J* (e.g., *n* = 200 and *J* = 100), ℬ_2_ becomes negligible compared to the variability of 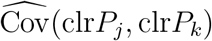. In other words, under the Sparse Dependence Assumption, the permutation replicates generated under the global null hypothesis, while preserving the zero-sum structure, serve as valid replicates for the observed data. Note that in each permutation, all clr*P*_*j*_ must be shuffled independently. If only clr*P*_*j*_ at taxon *j* is shuffled while all other taxa remain intact, the covariance after recenter-

ing becomes 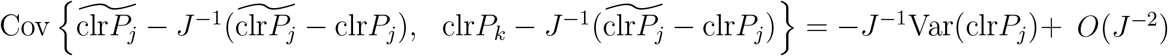, which does not align with the mean of the observed statistic, even under the Identity covariance structure where ℬ_2_ = 0. Moveover, if the mclr*P*_*j*_ values are shuffled, it becomes difficult to preserve the zero-sum structure at the clr*P*_*j*_ level, making it challenging to match the means of the statistics. Finally, we expect the means to align similarly for the nonlinear test statistics, although deriving a closed-form expression for their means is unattainable.

As done for the observed data, we modify the centered 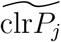 data at zero entries by replacing them by their taxon-specific means, where the zero statuses are tracked during the shuffling of clr*P*_*j*_ values. The centered and modified 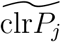 data are denoted by 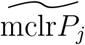, based on which we calculate the permutation statistics 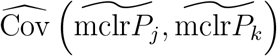 and d 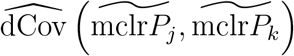. Finally, we obtain two-sided *p*-values for the linear tests and one-sided *p*-values for the nonlinear tests by comparing the permutation statistics with their observed counterparts.

The shuffling scheme can be readily extended to accommodate clustered data. To preserve the similarity of samples within the same subjects (or clusters), the clr*P*_*j*_ data from these samples should be shuffled as a whole across subjects and simultaneously shuffled within subjects. When subjects have unequal numbers of samples (i.e., cluster sizes), the shuffling should be stratified by subjects with the same cluster size.

We expect the linear test based on Pearson’s covariance to perform better when the dependence between a pair of taxa is linear, and the nonlinear test based on distance covariance to perform better when the dependence is nonlinear. Since we do not know the dependence type *a priori* for every pair of taxa, we construct an omnibus test for testing a *general* de-

pendence that combines the results from both tests for each pair. This omnibus test utilizes the existing permutation replicates. We use the minimum of the *p*-values obtained from the linear and nonlinear tests as the final test statistic and use the corresponding minima from the permutation replicates to simulate the null distribution [17], thereby obtaining the omnibus *p*-value.

Given the large number of *p*-values for all pairs of taxa from each type of test, linear, nonlinear or omnibus, we aim to declare correlated pairs by controlling the false discovery rate (FDR). Evidently, these *p*-values are not independent due to not only pairwise analysis but also taxon-taxon correlations. Fortunately, the Benjamini-Hochberg (BH) procedure [18], commonly used to control the FDR, allows for positive dependence among test statistics corresponding to null hypotheses. Positive dependence means that obtaining a significant result in one test increases the likelihood of obtaining another significant test. Our test statistics exhibit this dependence structure because obtaining a significant result for a pair of uncorrelated taxa, such as taxa A and B, increases the likelihood of obtaining a significant result for another pair of uncorrelated taxa, such as taxa A and C, if taxa B and C are correlated, either positively or negatively. Therefore, we apply the BH procedure to detect correlated pairs of taxa. In fact, we use Sandve’s procedure [19], a variation of BH that incorporates a sequential stopping rule for permutation-based multiple testing. This rule substantially improves computational efficiency by stopping the addition of replicates for tests once a pre-specified minimum number of extreme replicates has been reached or if the estimated FDR falls below a given threshold. The complete workflow of TestNet is depicted in Figure 8.

**Figure 8.**
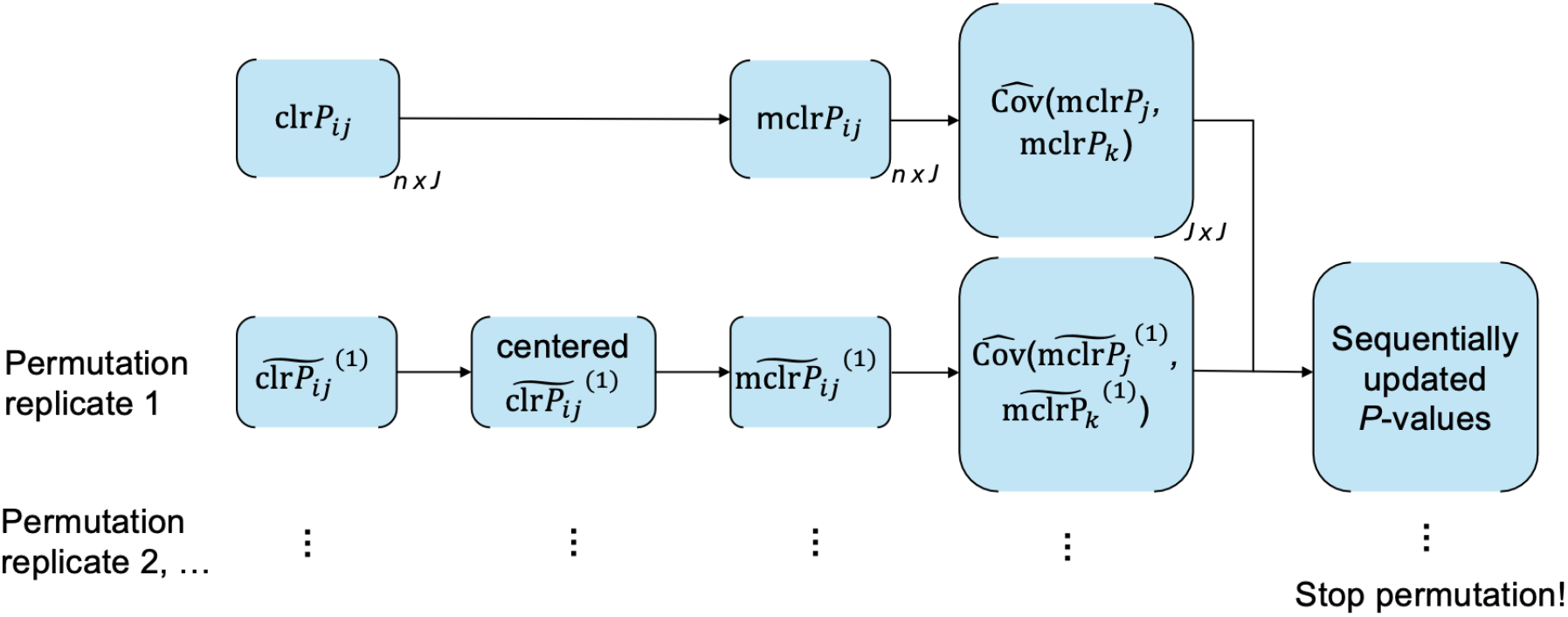
Workflow of TestNet, illustrated with the linear test. Each box represents a matrix. The superscript ‘(1)’ denotes the permutation replicate number.

### Simulation studies

To ensure a fair evaluation of all methods, we adopted the settings described in [9] and [10] to generate the underlying absolute abundance *Z*_*j*_ with various dependence structures. We considered the following seven covariance structures, **Σ** = {*σ*_*jk*_} = {Cov(log *Z*_*j*_, log *Z*_*k*_)}, for simulating linear dependencies, along with an additional dependence structure called U-shape for simulating nonlinear dependencies.

- (Identity) Set **Σ** = *I*_*J*_. It represents the global null and is an ideal case as ℬ_2_ = 0.
- (AR1+) Set *σ*_*jk*_ = 0.5 if |*j* − *k*| = 1, and otherwise 0. Set *σ*_*jj*_ = 1.
- (AR1) Similar to AR1+, but set *σ*_*jk*_ = ±0.5 if |*j* − *k*| = 1. The ± symbol indicates taking the positive and negative values with equal probabilities.
- (AR4) Set *σ*_*jk*_ = ±0.4 if |*j* − *k*| = 4, ±0.2 if |*j* − *k*| = 3 or 2, ±0.1 if |*j* − *k*| = 1, and otherwise 0. Set the diagonal elements (*σ*_*jj*_) large enough so that **Σ** is positive definite.
- (Sparse) Set **Σ** = diag(*A*_1_, *A*_2_), where 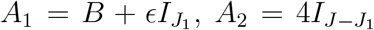, and 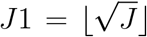. Here, *B* is a symmetric matrix with lower triangular entries sampled from the uniform distribution on [−1, −0.5] ∪ [0.5, 1] with probability 0.3, and otherwise 0. The term *ϵ* = max(−*λ*_min_(*B*), 0)+0.01, where *λ*_min_(*B*) denotes the smallest eigenvalue of *B*, ensures that *A*_1_ is positive definite.
- (Hub) Sample three hub taxa from the top 20 most abundant taxa. For a hub taxon *j* and any other taxon *k*, set *σ*_*jk*_ = ±0.3 with probability 0.7, and otherwise 0. For two non-hub taxa *j* and *k*, set *σ*_*jk*_ = ±0.3 with probability 0.2, and otherwise 0. Set *σ*_*jj*_s large enough so that **Σ** is positive definite.
- (Block) Divide the *J* taxa equally into 10 blocks. For each pair (*j, k*) within a block, set *σ*_*jk*_ = ±0.3 with probability 0.5, and otherwise 0. For each pair (*j, k*) between blocks, set *σ*_*jk*_ = ±0.3 with probability 0.2, and otherwise 0. Set *σ*_*jj*_s large enough so that **Σ** is positive definite.
- (U-shape) Group every two taxa together. Set a nonlinear, U-shaped correlation for the two taxa within each group, while keeping the taxa in different groups independent.

In the linear dependence settings, we generated the log *Z*_*j*_ data with a given variance-covariance matrix **Σ** and a mean vector *µ* (each element of *µ* sampled from *U* [0, 10]) from either the Normal or Gamma distribution. For the Gamma distribution, we set log 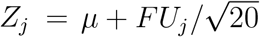, where *FF* ^T^ = **Σ** and the components of *U*_j_ are independent gamma variables with a shape parameter of 10 and a scale parameter of 1. In the U-shape dependence setting, we first generated the log *Z*_*j*_ data independently for one taxon in each group, and then generated the log *Z*_*j*_ data for the other taxon in the same group as the orthogonal polynomial of degree 2 from the log *Z*_*j*_ data of the first taxon, plus an error term drawn from *N* (0, 0.05). The underlying relative abundance were calculated as 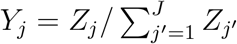.

In some cases, we simulated clustered data with four samples in each cluster. We generated each cluster-specific mean as *µ*_*i*_ = *µ* + *ϵ*_*i*_, where *ϵ*_*i*_ ~ *N* (0, 1), and assigned this mean to all samples within the cluster. In other cases, we introduced experimental bias by calculating 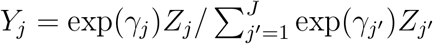, where the bias factors *γ*_j_ s were sampled from *N* (0, 0.5).

Finally, we generated the read count data *X*_*j*_ independently for each taxon, given *Y*_*j*_ and the library size *l*, from the Negative-Binomial distribution with a mean of *lY*_*j*_ and an overdispersion parameter of 1. Here, *l* was sampled from *N* (10000, (10000*/*3)^2^) and left-truncated at 2000. This overdispersion value produced realistic proportions of zeros, similar to those observed in real microbiome data. The Dirichlet-Multinomial distribution was not used because it leads to negative correlations across taxa even when *Y*_*j*_ is fixed. It is important to note that this step of generating *X*_*j*_ was often missed in simulation studies conducted for existing methods such as COAT and CCLasso [8, 9], where *Y*_*j*_ was directly used as the observed relative abundance *P*_*j*_ to evaluate those methods. Consequently, those evaluations ignored the effect of sequencing variation.

Throughout this work, the number of taxa *J* was set to 100 unless otherwise specified as 50. We considered these relatively small *J* values because our method performs better with larger *J* values. Prior to analysis, it is customary to filter out very rare taxa that are likely contaminants and increase the burden of multiple testing. We considered two filtering criteria. The stringent criterion filters out taxa with mean relative abundances less than 0.1%, which tends to reduce the number of taxa by 50%. The lenient criterion filters out taxa that are present in less than 5% of all samples, which often led to no exclusions in our simulated data. The stringent criterion was used as the default unless otherwise specified.

### Verifying the Sparse Dependence Assumption

Figure S15 compares the two sides of the inequality in the Sparse Dependence Assumption, demonstrating that the assumption holds across all scenarios considered. The x-axis represents the sample variance of clr*P*_*j*_, calculated from read count data based on *n* = 10000 samples, a large sample size to ensure accurate variance estimation.

### Examining the bias terms ℬ_1_– ℬ_4_

Figure 9 compares the sample variances and covariances of log *Z*_*j*_, clr*Z*_*j*_ (equivalently, clr*Y*_*j*_), and clr*P*_*j*_, based on *n* = 10000 samples. The upper panel shows that 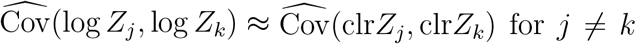 and 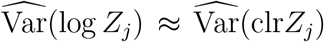, indicating that the bias ℬ_1_ + ℬ_2_ is small, though noticeable in some cases such as AR1+. The middle panel shows that 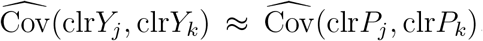, suggesting that ℬ_3_ is small. However,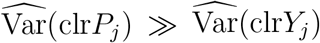 in most cases, indicating that ℬ_4_ is large. The lower panel shows that 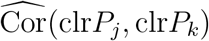 is significantly attenuated compared to 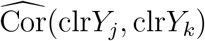. Distance variances and covariances for measuring nonlinear dependencies are displayed in Figure S16, showing similar comparisonresults, with the exception that 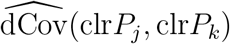 tends to be smaller than 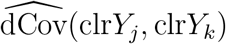.

**Figure 9.**
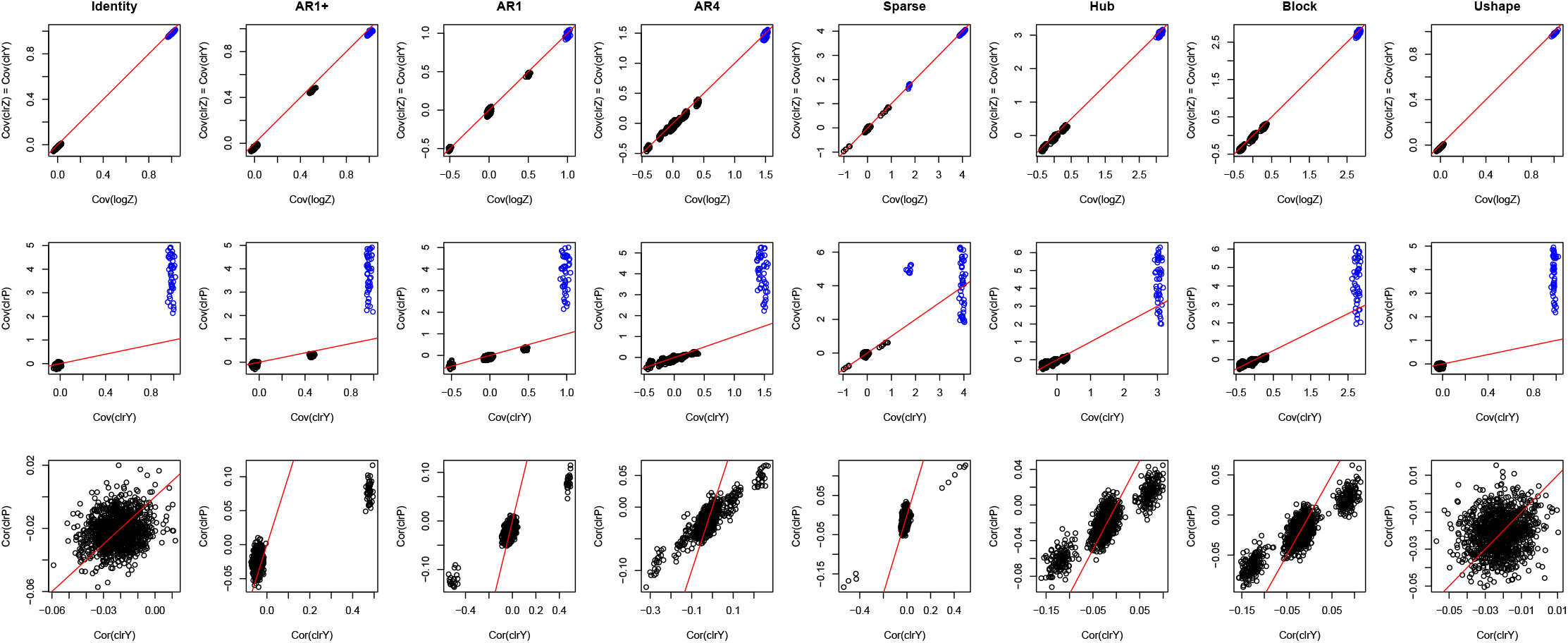
Upper panel: Sample Pearson’s covariances (in black) and variances (in blue) of log *Z*_*j*_ and clr*Z*_*j*_. Middle panel: Sample Pearson’s covariances (in black) and variances (in blue) of clr*Y*_*j*_ and clr*P*_*j*_. Lower panel: Sample Pearson’s correlations of clr*Y*_*j*_ and clr*P*_*j*_. The red line represents the 45^°^ reference line.

### Evaluating the modification of clr*P*_*j*_ at zero entries

Figure S17 displays the clr*P*_*j*_ values before and after the modification of clr*P*_*j*_ at zero entries for two pairs of taxa, based on *n* = 1000 samples. The upper panel demonstrates that for a pair of truly uncorrelated taxa, both of which have many zeros, the clr*P*_*j*_ values show a spurious Pearson’s correlation of 0.45, which reduces to merely −0.013 after the modification. The lower panel indicates that for a pair of truly correlated taxa, the modification has a minimal effect on the correlation measure, which is 0.14 before and 0.15 after modification.

### Assessing the validity of permutation replicates

Figure S18 compares the distributions of observed and permutation test statistics across all null taxon pairs based on one replicate of observed data and one permutation replicate. For inferring linear dependencies (upper panel), the observed statistics (i.e., sample Pearson’s covariances) show a slight shift to the left of zero with *n* = 100 samples, which becomes more evident with *n* = 1000 samples. However, the corresponding permutation statistics without centering 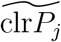 are centered at zero, creating a discrepancy from the observed statistics. After centering 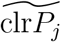, the distribution of the permutation statistics matches well with that of the observed statistics. For inferring nonlinear dependencies (lower panel), the match between the observed statistics (i.e., sample distance covariances) and the corresponding permutation statistics also occurs after centering 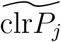. This match was verified for all dependence structures and with both the stringent and lenient filters, as shown in Figures S19–S22.

## Acknowledgement

Not applicable.

## Availability of data and materials

The R package `TestNet` and simulation code are available on GitHub at https://github.com/yijuanhu/TestNet. The rectal mucosal microbiome dataset and the gut microbiome dataset are both available in the “data” folder.

## Authors’ contributions

Y.-J.H. conceived this work, developed methodologies, conducted numerical studies, and wrote the manuscript; C.S. developed methodologies, conducted numerical studies and revised the manuscript and revised the manuscript; M.H. conducted numerical studies; V.E.V.D. and C.F.K. interpreted results and revised the manuscript.

## Consent for publication

No consent for publication was required for this study; all microbiome datasets used here are publicly available.

## Ethics approval and consent to participate

This study only involved secondary analyses of existing, de-identified datasets; as such it does not require separate IRB consent.

## References

1. Banerjee S, Schlaeppi K, van der Heijden MG. Keystone taxa as drivers of microbiome structure and functioning. Nature Reviews Microbiology. 2018;16(9):567–576.

2. Wu G, Zhao N, Zhang C, Lam YY, Zhao L. Guild-based analysis for understanding gut microbiome in human health and diseases. Genome medicine. 2021;13:1–12.

3. Aitchison J. The statistical analysis of compositional data. Chapman and Hall, London-New York; 1986.

4. Gloor GB, Macklaim JM, Pawlowsky-Glahn V, Egozcue JJ. Microbiome datasets are compositional: and this is not optional. Frontiers in microbiology. 2017;8:2224.

5. Mandal S, Van Treuren W, White RA, Eggesbø M, Knight R, Peddada SD. Analysis of composition of microbiomes: a novel method for studying microbial composition. Microbial ecology in health and disease. 2015;26(1):27663.

6. Hu Y, Satten GA, Hu YJ. LOCOM: A logistic regression model for testing differential abundance in compositional microbiome data with false discovery rate control. Proceedings of the National Academy of Sciences. 2022;119(30):e2122788119.

7. Friedman J, Alm EJ. Inferring correlation networks from genomic survey data. PLoS computational biology. 2012;8(9):e1002687.

8. Fang H, Huang C, Zhao H, Deng M. CCLasso: correlation inference for compositional data through Lasso. Bioinformatics. 2015;31(19):3172–3180.

9. Cao Y, Lin W, Li H. Large covariance estimation for compositional data via composition-adjusted thresholding. Journal of the American Statistical Association. 2019;114(526):759–772.

10. Lin H, Eggesbø M, Peddada SD. Linear and nonlinear correlation estimators unveil undescribed taxa interactions in microbiome data. Nature Communications. 2022;13(1):4946.

11. McLaren MR, Willis AD, Callahan BJ. Consistent and correctable bias in metagenomic sequencing experiments. Elife. 2019;8:e46923.

12. Zhu Z, Satten GA, Mitchell C, Hu YJ. Constraining PERMANOVA and LDM to within-set comparisons by projection improves the efficiency of analyses of matched sets of microbiome data. Microbiome. 2021;9(1):1–19.

13. Lin H, Peddada S. Multi-group Analysis of Compositions of Microbiomes with Covariate Adjustments and Repeated Measures. Nature methods. 2024;21:83–91.

14. Kurtz ZD, Müller CL, Miraldi ER, Littman DR, Blaser MJ, Bonneau RA. Sparse and compositionally robust inference of microbial ecological networks. PLoS computational biology. 2015;11(5):e1004226.

15. Székely GJ, Rizzo ML, Bakirov NK. Measuring and testing dependence by correlation of distances. The Annals of Statistics. 2007;35(6):2769—-2794.

16. Székely GJ, Rizzo ML. Energy statistics: A class of statistics based on distances. Journal of statistical planning and inference. 2013;143(8):1249–1272.

17. Westfall PH, Young SS. Resampling-based multiple testing: Examples and methods for p-value adjustment. John Wiley & Sons; 1993.

18. Benjamini Y, Hochberg Y. Controlling the false discovery rate: a practical and powerful approach to multiple testing. Journal of the royal statistical society Series B (Methodological). 1995;p. 289–300.

19. Sandve GK, Ferkingstad E, Nygård S. Sequential Monte Carlo multiple testing. Bioinformatics. 2011;27(23):3235–3241.

20. Wang J, Kurilshikov A, Radjabzadeh D, Turpin W, Croitoru K, Bonder MJ, et al. Meta-analysis of human genome-microbiome association studies: the MiBioGen consortium initiative. Springer; 2018.

21. Markowitz RHG, LaBella AL, Shi M, Rokas A, Capra JA, Ferguson JF, et al. Microbiome-associated human genetic variants impact phenome-wide disease risk. Proceedings of the National Academy of Sciences. 2022;119(26):e2200551119.

22. Van Doren VE, Ackerley CG, Arthur RA, Murray PM, Smith SA, Hu YJ, et al. Rectal mucosal inflammation, microbiome, and wound healing in men who have sex with men who engage in receptive anal intercourse. Scientific Report. 2024;14:31598.

23. Ackerley CG, Smith SA, Murray PM, Amancha PK, Arthur RA, Zhu Z, et al. The rectal mucosal immune environment and HIV susceptibility among young men who have sex with men. Frontiers in Immunology. 2022;13:972170.

24. Wu GD, Chen J, Hoffmann C, Bittinger K, Chen YY, Keilbaugh SA, et al. Linking long-term dietary patterns with gut microbial enterotypes. Science. 2011;334(6052):105–108.

