## Supplemental materials for "TestNet: A Testing Method for Inferring Microbial Networks with False Discovery Rate Control, for Clustered and Unclustered Samples"

### Supplementary Materials

#### Text A: Approximation of the bias term $\mathcal{B}_3$

Note that $\mathrm{clr}P_j = \mathrm{clr}(X_j + 0.5)$. For simplicity, we assume that $X_j$ here and in Text B already includes the pseudo count 0.5. Let $\overline{\log X} = J^{-1}\sum_{j'} \log X_{j'}$. Under the Independence Assumption, the left-hand side of (1) expands to

$$\begin{aligned}&\mathrm{E}\left[\mathrm{Cov}(\log X_j - \overline{\log X}, \log X_k - \overline{\log X}|\mathbb{Y})\right] \\ &\qquad = -J^{-1}\mathrm{E}\left[\mathrm{Var}(\log X_j|\mathbb{Y})\right] - J^{-1}\mathrm{E}\left[\mathrm{Var}(\log X_k|\mathbb{Y})\right] + J^{-2}\sum_{j'}\mathrm{E}\left[\mathrm{Var}(\log X_{j'}|\mathbb{Y})\right].\end{aligned}$$

On the right-hand side of (1), each variance term $\mathrm{E}\left[\mathrm{Var}(\mathrm{clr}P_j|\mathbb{Y})\right]$ expands to

$$\mathrm{E}\left[\mathrm{Var}(\log X_j - \overline{\log X}|\mathbb{Y})\right] = \mathrm{E}\left[\mathrm{Var}(\log X_j|\mathbb{Y})\right] - 2J^{-1}\mathrm{E}\left[\mathrm{Var}(\log X_j|\mathbb{Y})\right] + J^{-2}\sum_{j'}\mathrm{E}\left[\mathrm{Var}(\log X_{j'}|\mathbb{Y})\right],$$

which is essentially $\mathrm{E}\left[\mathrm{Var}(\log X_j|\mathbb{Y})\right] + O(J^{-1})$. Therefore, the two sides of (1) are approximately equal.

#### Text B: The bias term $\mathcal{B}_4$ due to overdispersion

Using the delta method, we obtain

$$\mathrm{Var}(\mathrm{clr}X_j|\mathbb{Y}) \approx \left(\frac{\partial \mathrm{clr}X_j}{\partial \mathbb{X}}\right)_{lY_j}^{\mathrm{T}} \mathrm{Var}(\mathbb{X}|\mathbb{Y}) \left(\frac{\partial \mathrm{clr}X_j}{\partial \mathbb{X}}\right)_{lY_j},$$

where $(\partial \mathrm{clr}X_j/\partial \mathbb{X})_{lY_j}$ is the derivative of $\mathrm{clr}X_j$ with respect to $\mathbb{X} \equiv (X_1, X_2, \ldots, X_J)$, evaluated at $\mathrm{E}(X_j|\mathbb{Y}) = lY_j$, and is given by

$$\left\{-(JlY_1)^{-1}, \ldots, (J-1)(JlY_j)^{-1}, \ldots, -(JlY_J)^{-1}\right\}^{\mathrm{T}}.$$

Under the Independence Assumption and for sufficiently large $J$, $\mathrm{Var}(\mathrm{clr}X_j|\mathbb{Y})$ is calculated as approximately $l^{-2}Y_j^{-2}\mathrm{Var}(X_j|\mathbb{Y})$. Therefore, In the presence of overdispersion that $\mathrm{Var}(X_j|\mathbb{Y})$ is at least $O(l^2)$, $\mathcal{B}_4$ remains significant even as $l$ approaches infinity.

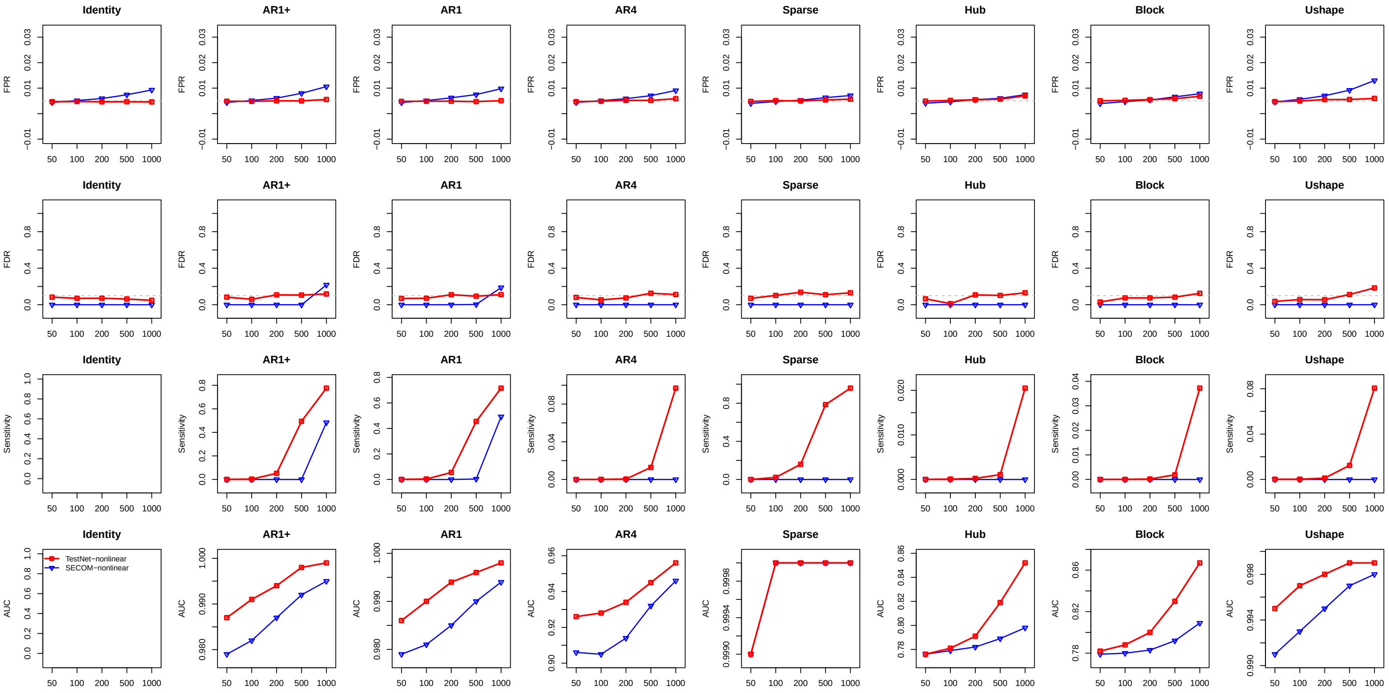


Figure S1: Results on inference of nonlinear dependencies based on data filtered with the lenient criterion. Refer to the caption of Figure 1 for additional information.

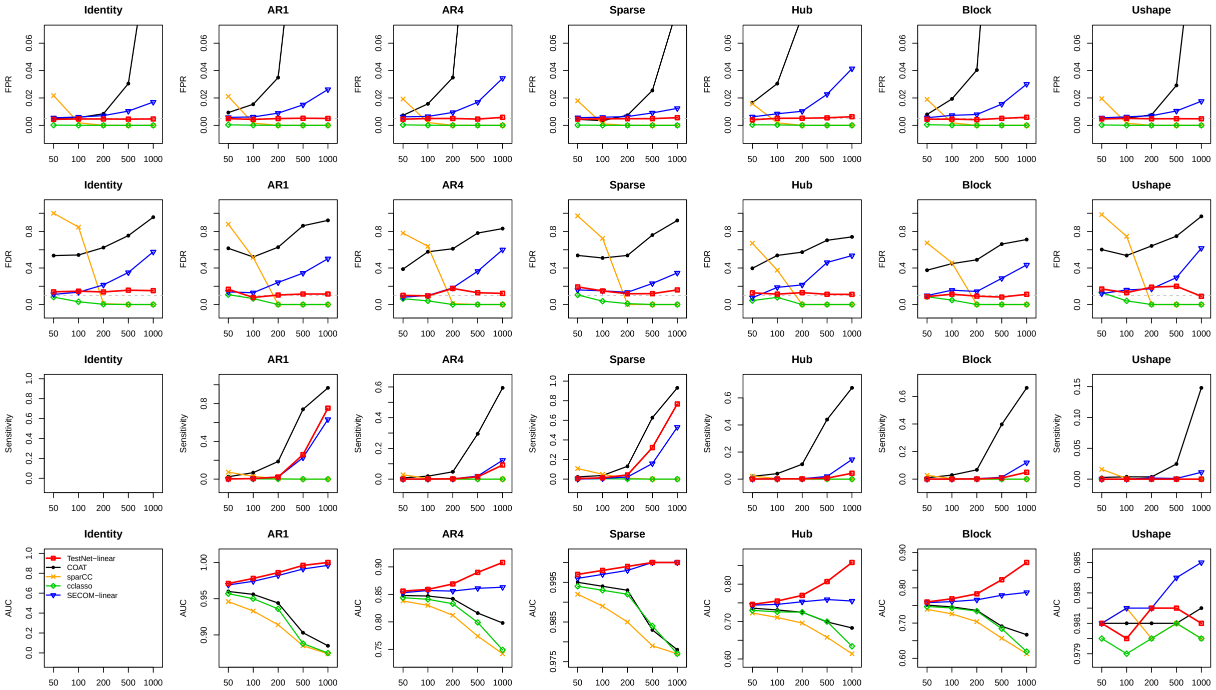

Figure S2: Results on inference of linear dependencies based on data simulated with Gamma distributions and filtered with the stringent criterion. The AR1+ structure led to many simulated replicates of data with only one taxon in the community and thus omitted here. Refer to the caption of Figure 1 for additional information.

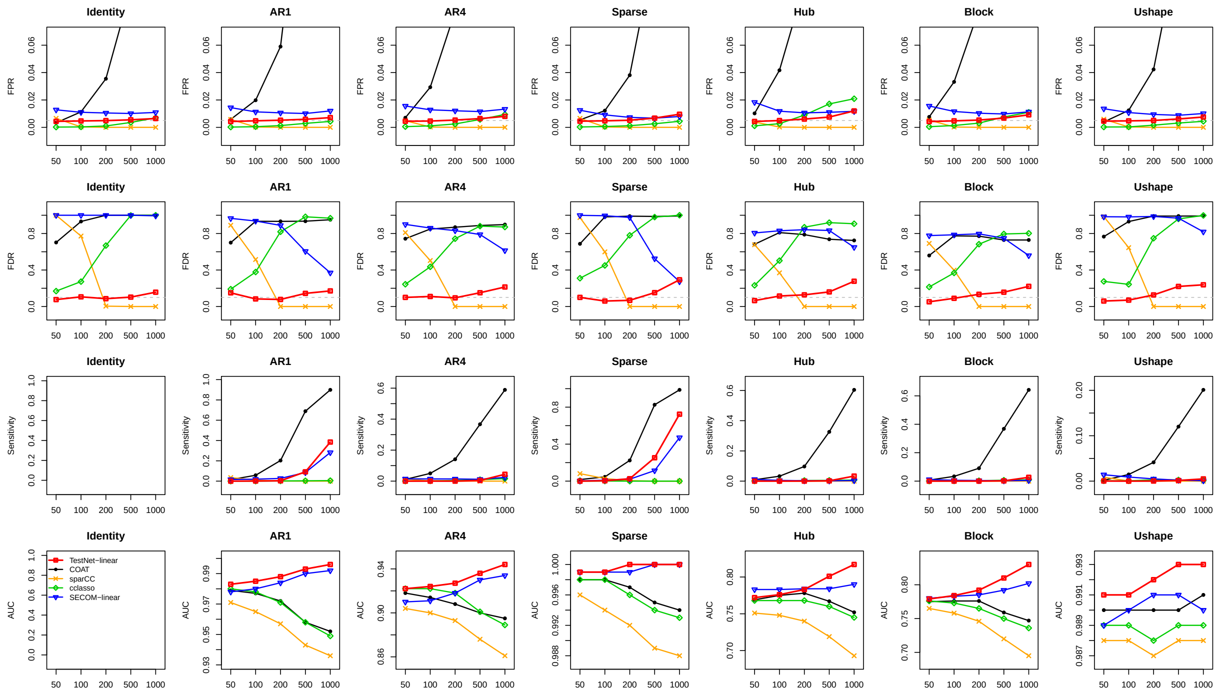


Figure S3: Results on inference of linear dependencies based on data simulated with Gamma distributions and filtered with the lenient criterion. Refer to the caption of Figure 1 for additional information.

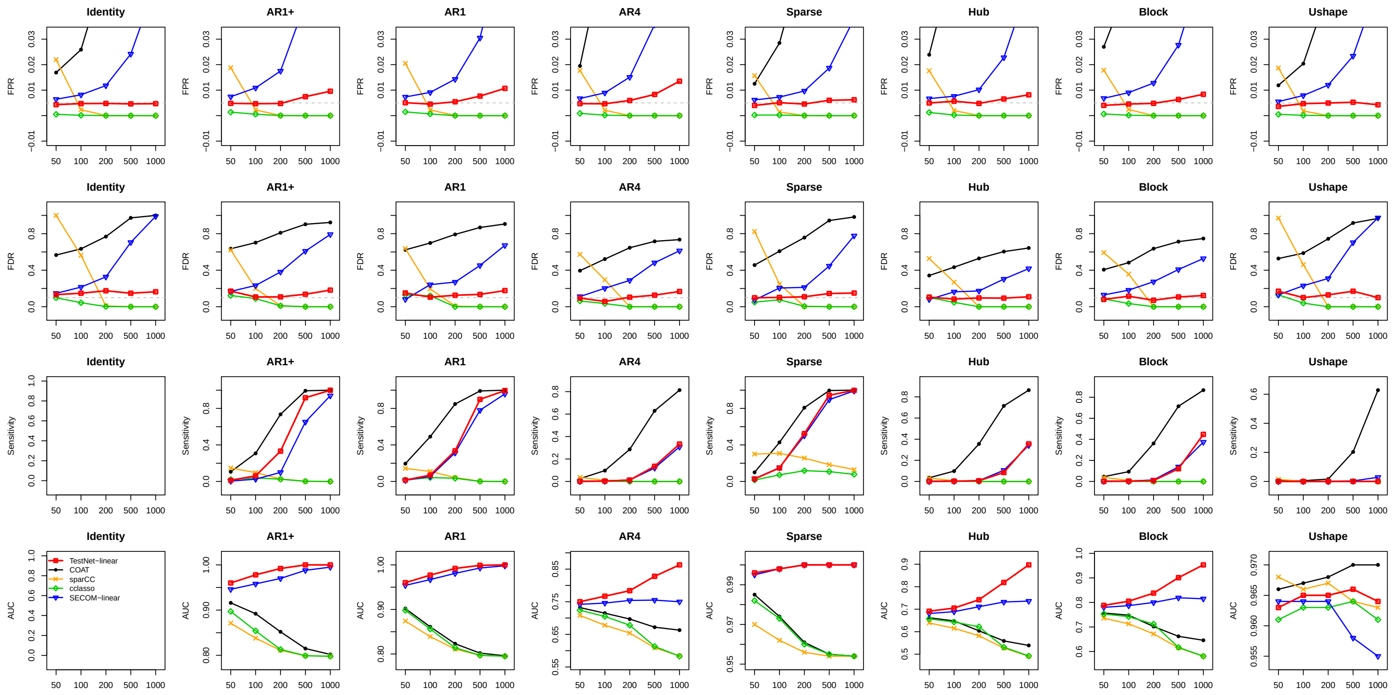

Figure S4: Results on inference of linear dependencies based on data simulated with $J = 50$ and filtered with the stringent criterion. Refer to the caption of Figure 1 for additional information.

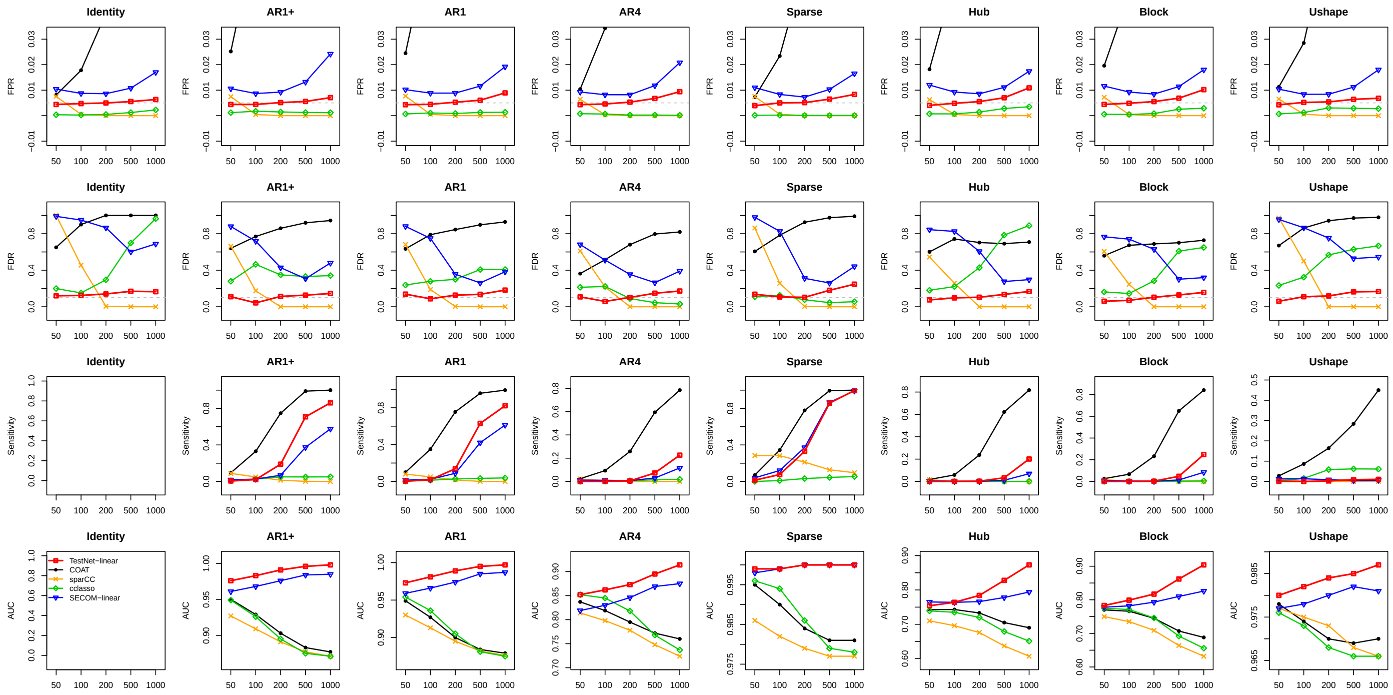


Figure S5: Results on inference of linear dependencies based on data simulated with $J = 50$ and filtered with the lenient criterion. Refer to the caption of Figure 1 for additional information.

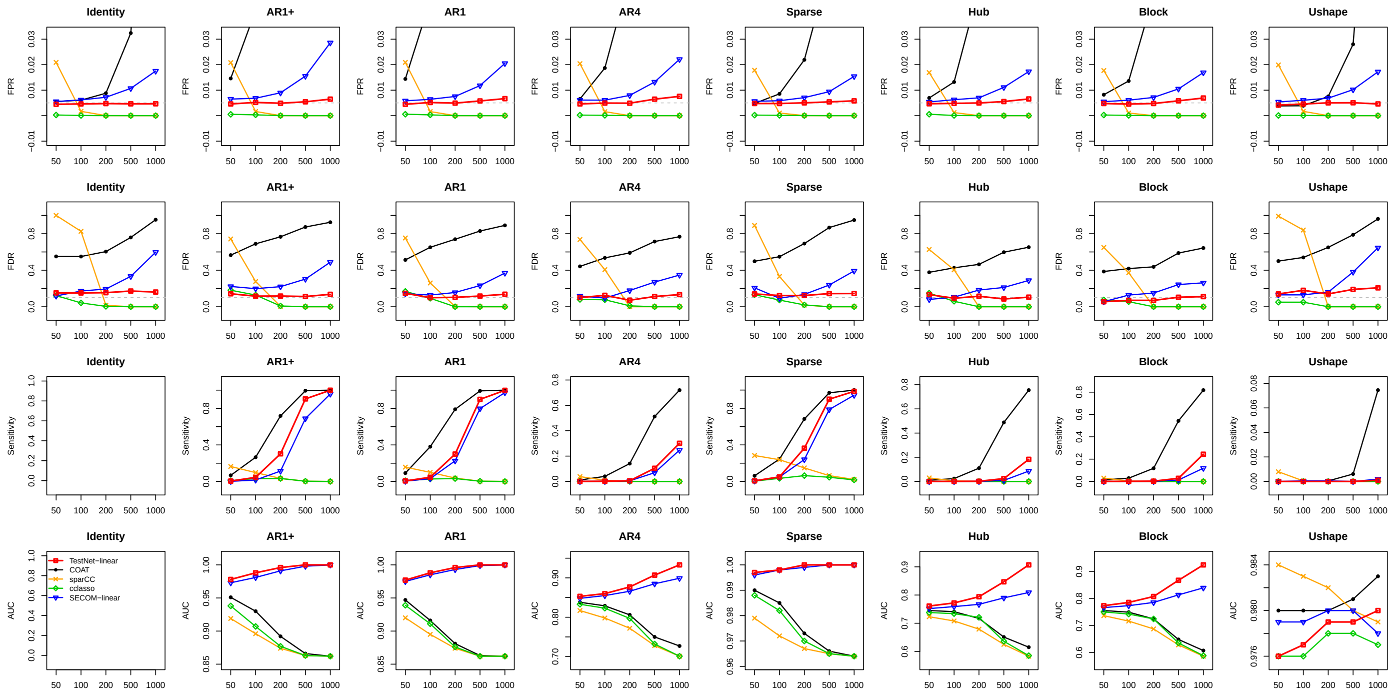


Figure S6: Results on inference of linear dependencies based on data simulated with experimental bias and filtered with the stringent criterion. Refer to the caption of Figure 1 for additional information.

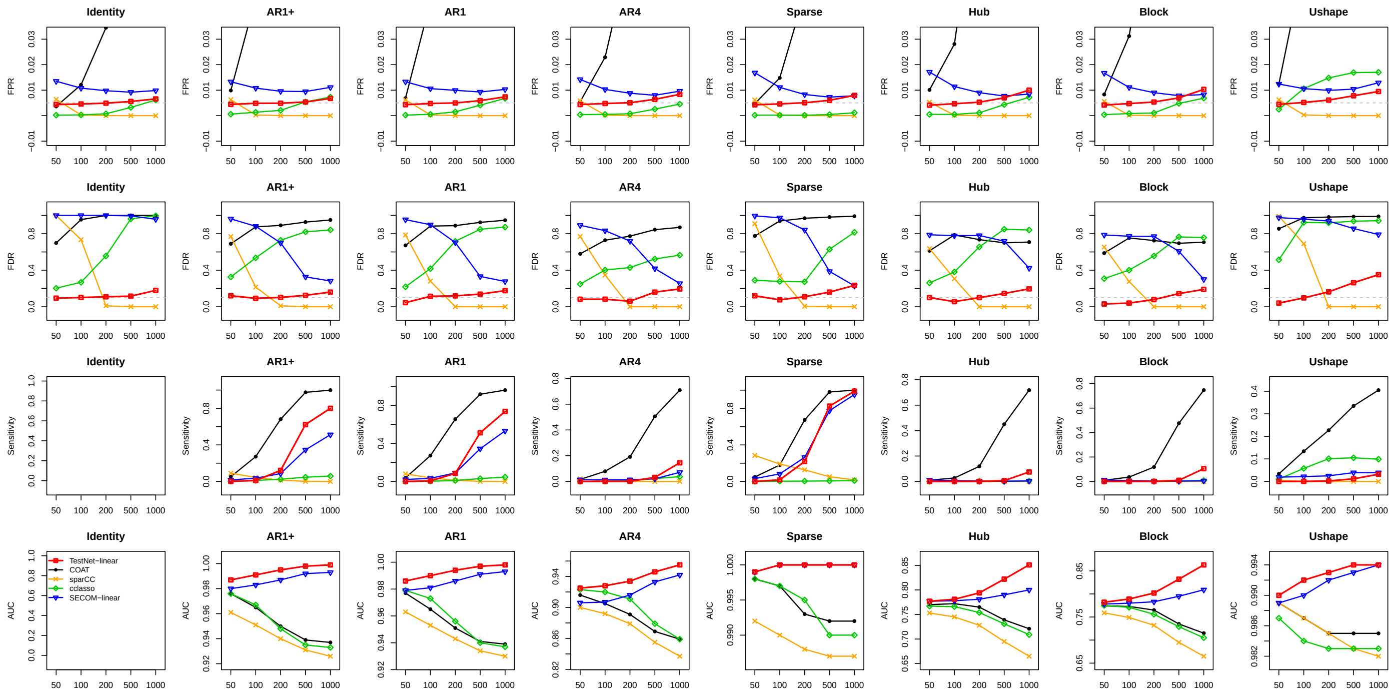


Figure S7: Results on inference of linear dependencies based on data simulated with experimental bias and filtered with the lenient criterion. Refer to the caption of Figure 1 for additional information.

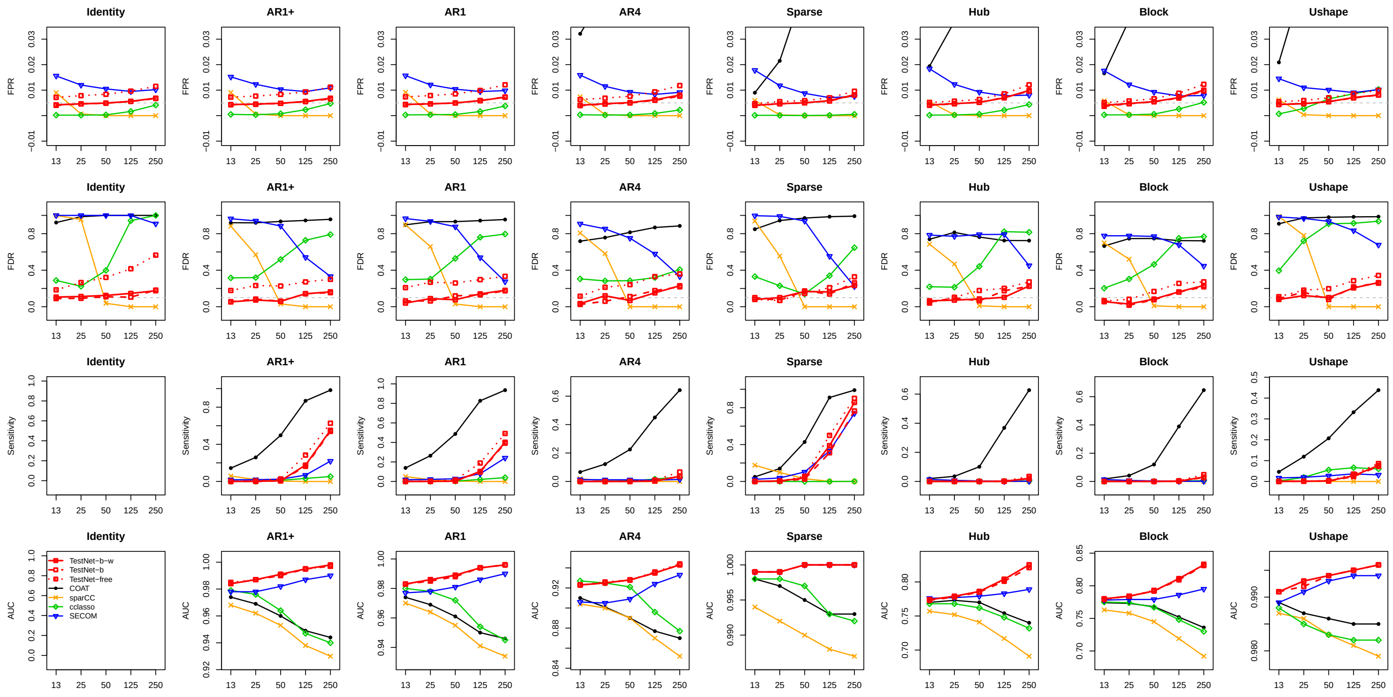

Figure S8: Results on inference of linear dependencies based on clustered data filtered with the lenient criterion. Refer to the caption of Figure 4 for additional information.

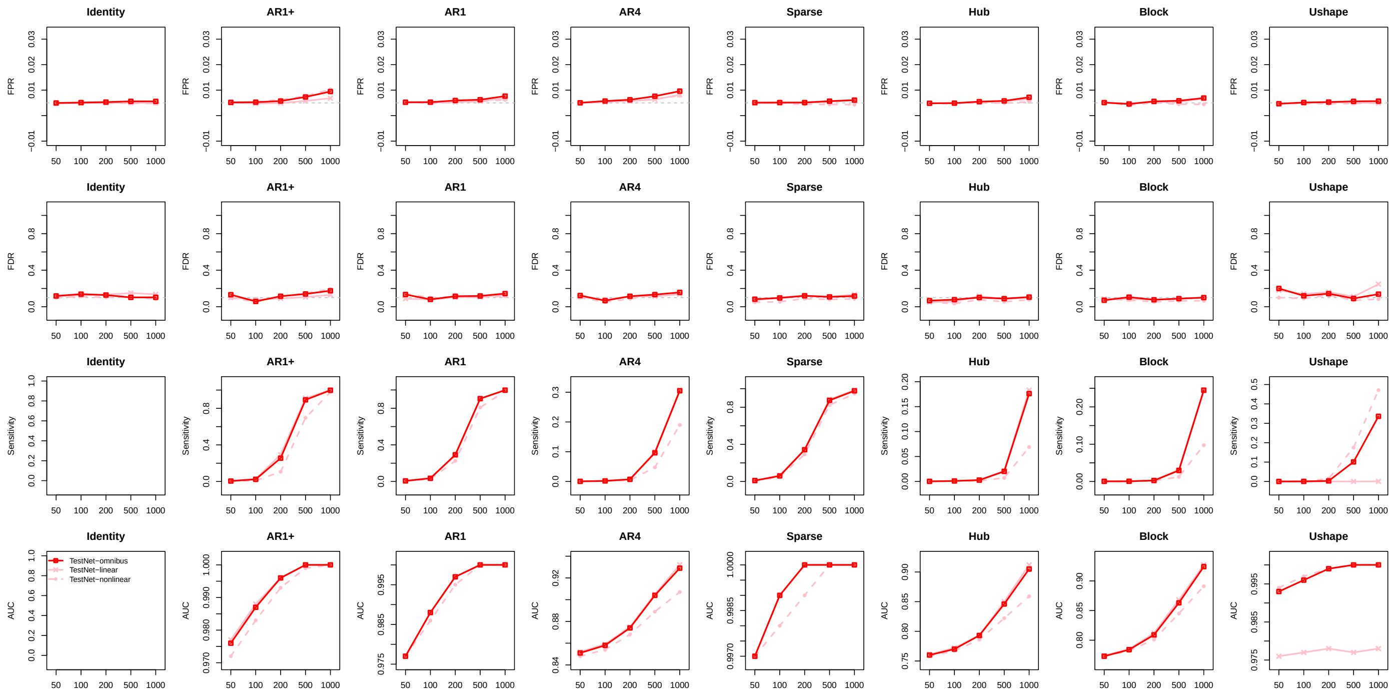


Figure S9: Results on inference of linear, nonlinear and general dependencies by TestNet, based on data filtered with the stringent criterion. Refer to the caption of Figure 1 for additional information.

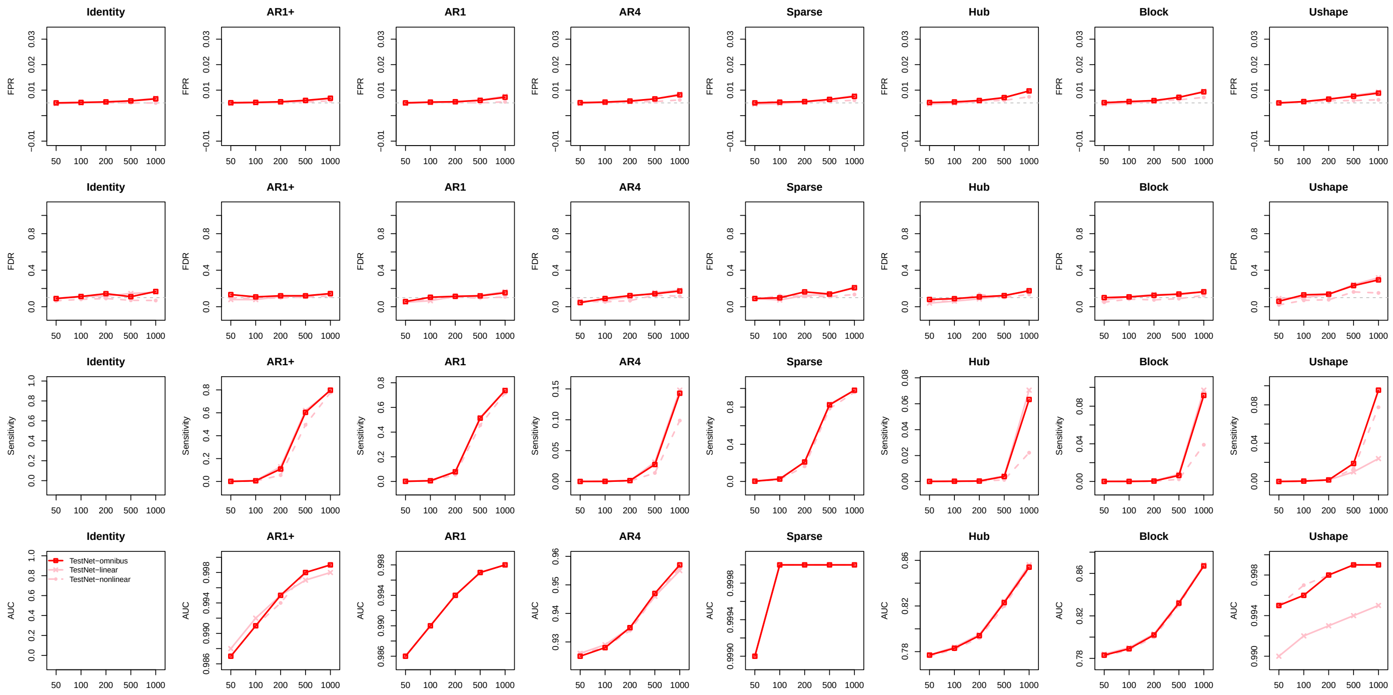

Figure S10: Results on inference of linear, nonlinear and general dependencies by TestNet, based on data filtered with the lenient criterion. Refer to the caption of Figure 1 for additional information.

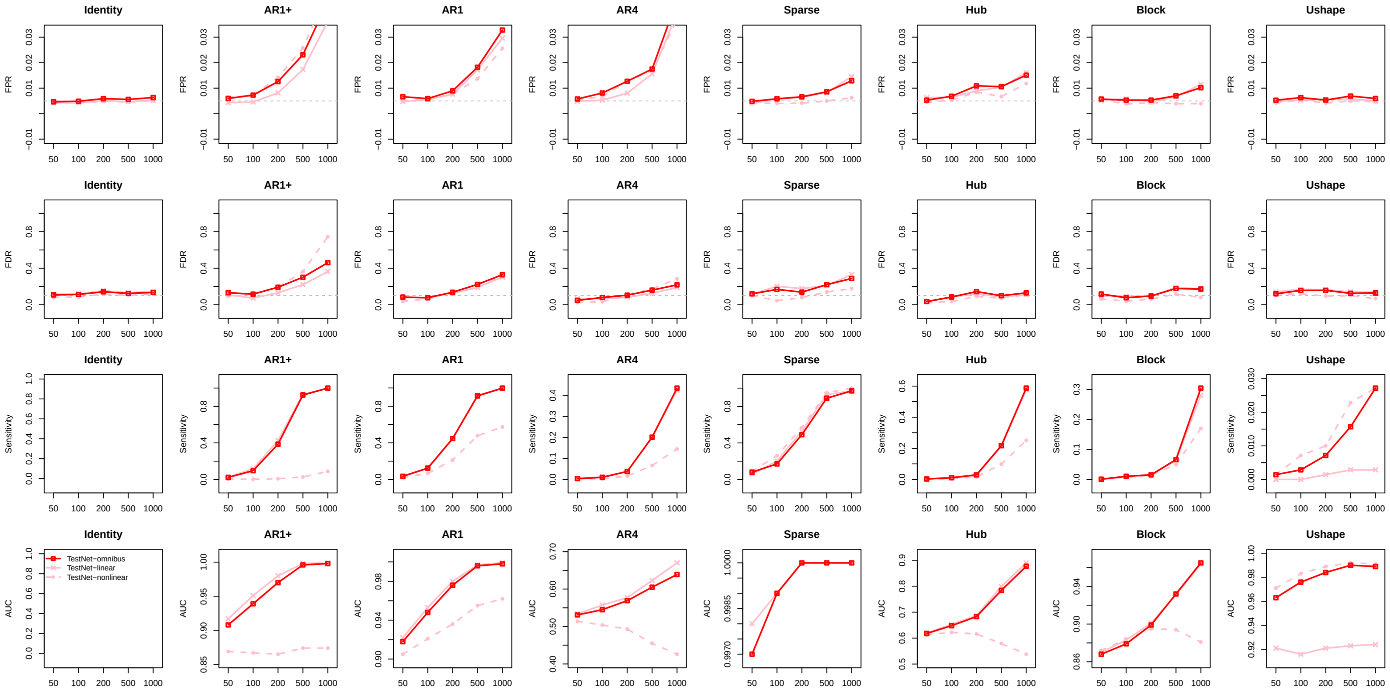

Figure S11: Results on inference of linear, nonlinear and general dependencies by TestNet, based on data simulated with $J = 20$ and filtered with the stringent criterion. Refer to the caption of Figure 1 for additional information.

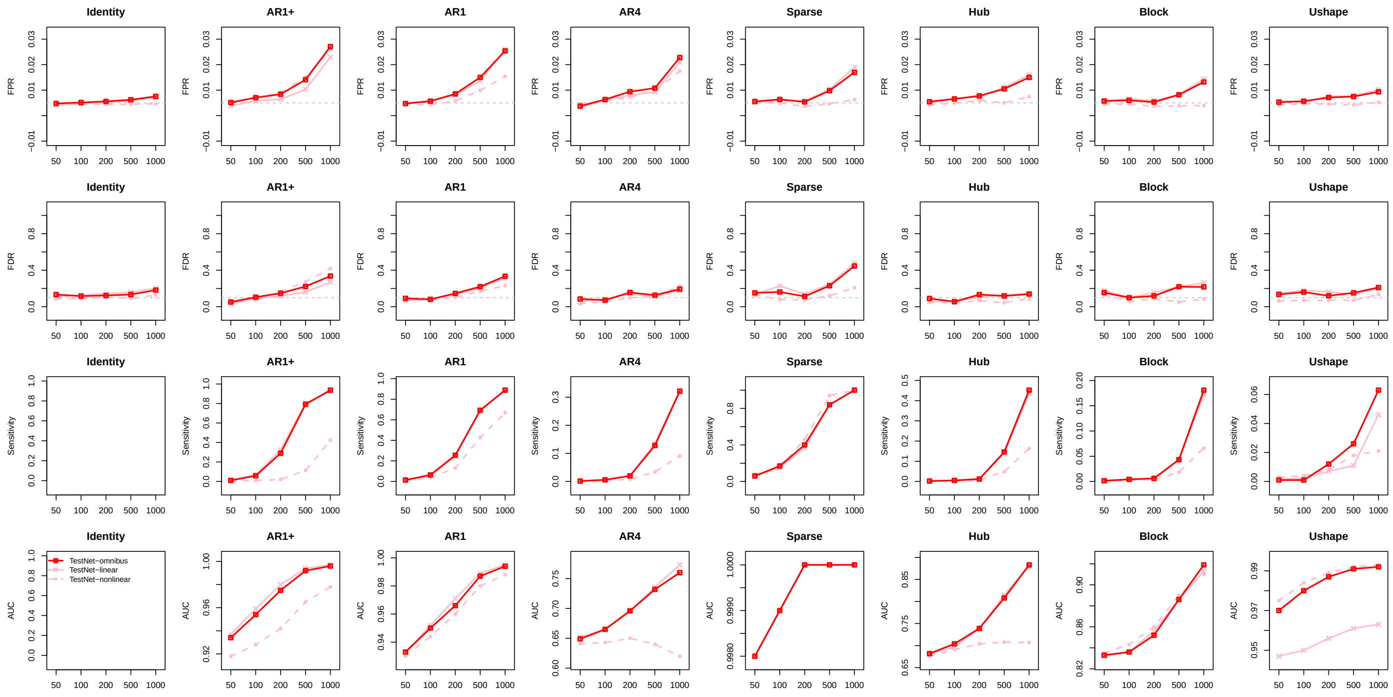

Figure S12: Results on inference of linear, nonlinear and general dependencies by TestNet, based on data simulated with $J = 20$ and filtered with the lenient criterion. Refer to the caption of Figure 1 for additional information.

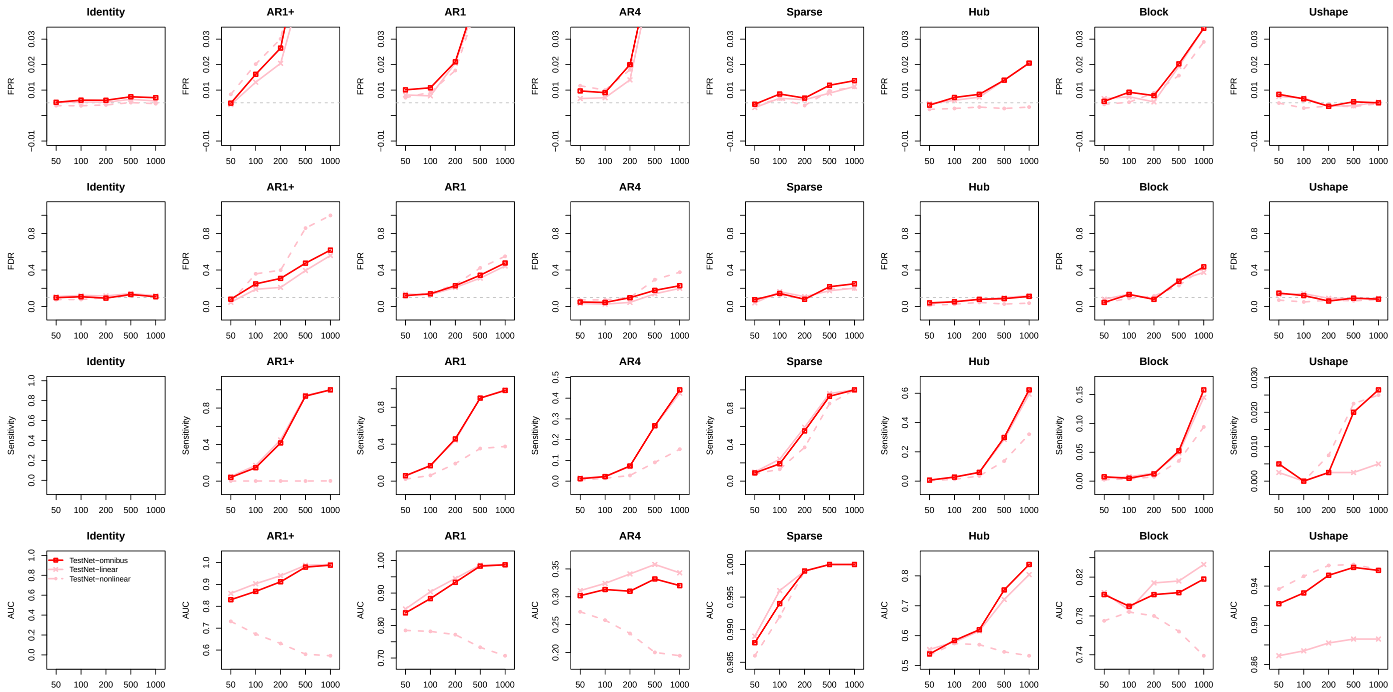

Figure S13: Results on inference of linear, nonlinear and general dependencies by TestNet, based on data simulated with $J = 10$ and filtered with the stringent criterion. Refer to the caption of Figure 1 for additional information.

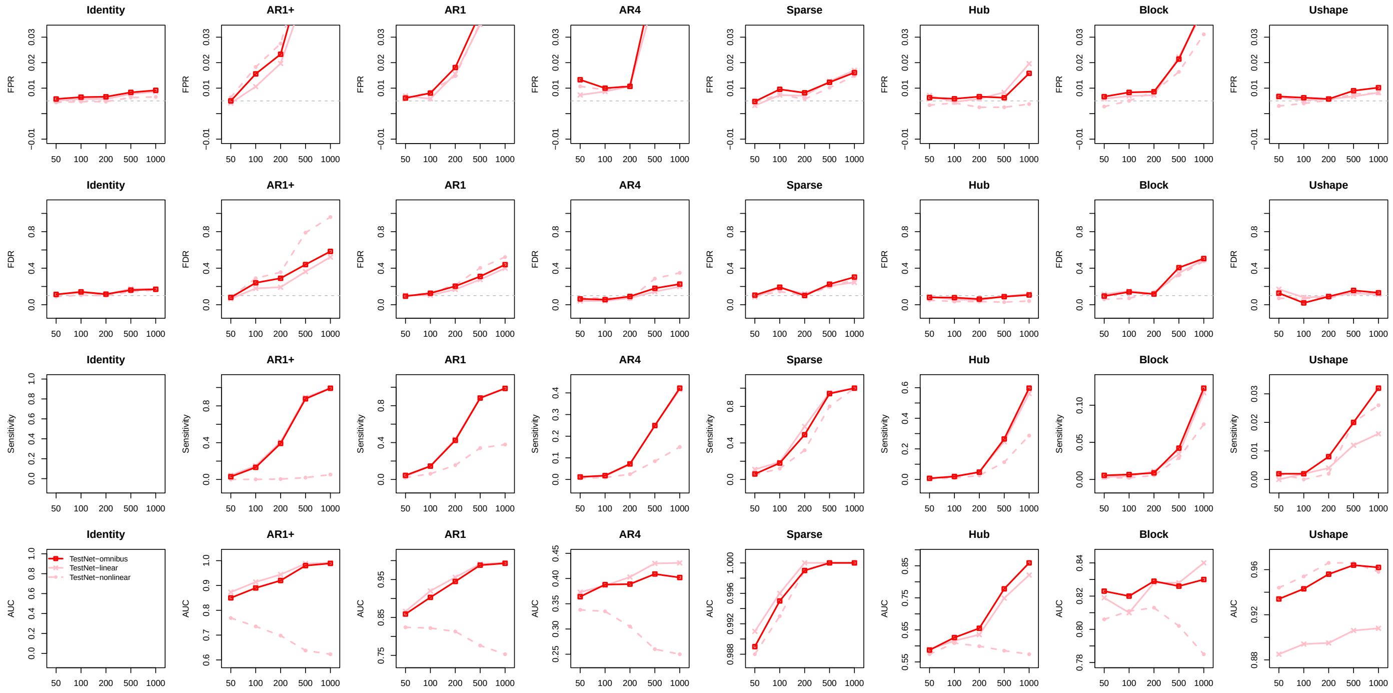


Figure S14: Results on inference of linear, nonlinear and general dependencies by TestNet, based on data simulated with $J = 10$ and filtered with the lenient criterion. Refer to the caption of Figure 1 for additional information.

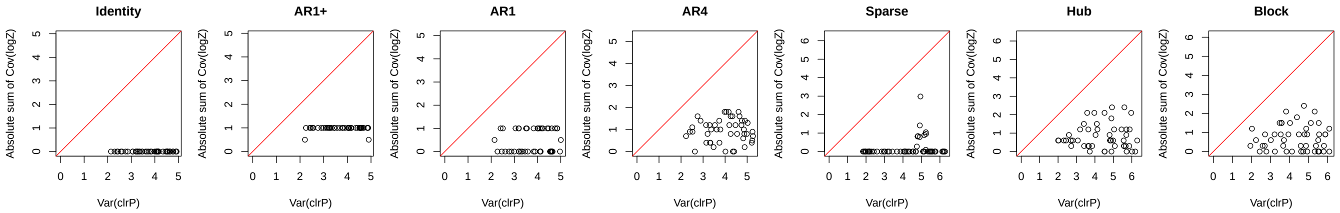


Figure S15: Comparing the left-hand side (y-axis) and the right-hand side (x-axis) of the inequality in the Sparse Dependence Assumption.

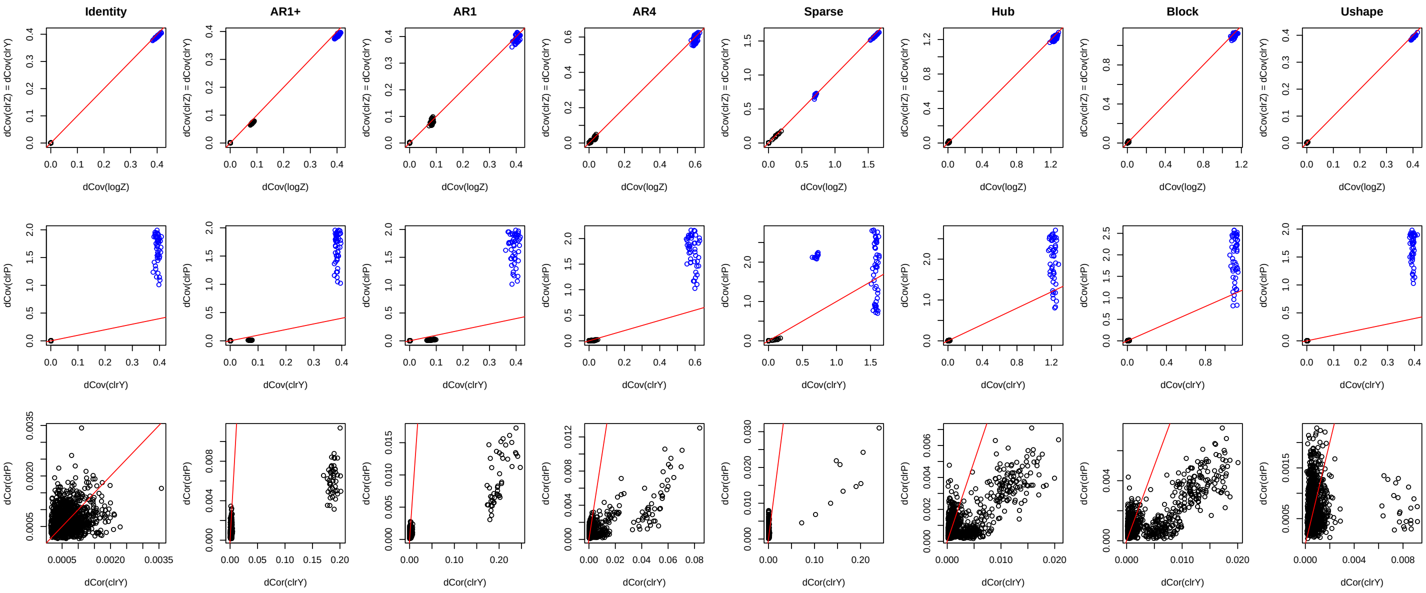


Figure S16: Upper panel: Sample distance covariances (in black) and variances (in blue) of $\log Z_j$ and $\mathrm{clr} Z_j$. Middle panel: Sample distance covariances (in black) and variances (in blue) of $\mathrm{clr} Y_j$ and $\mathrm{clr} P_j$. Lower panel: Sample distance correlations of $\mathrm{clr} Y_j$ and $\mathrm{clr} P_j$. The red line represents the 45° reference line. This is the nonlinear counterpart of Figure 9.

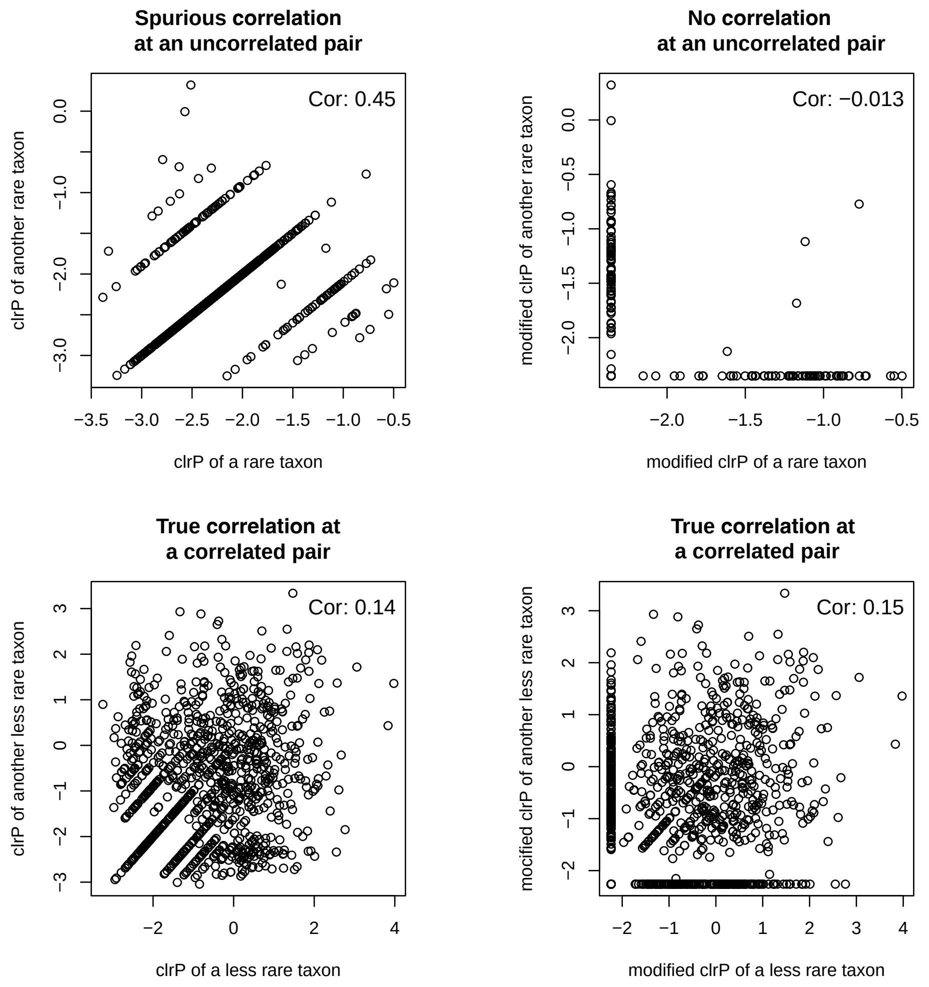


Figure S17: The clr$P_j$ values for an uncorrelated pair of taxa (upper panel) and a correlated pair of taxa (lower panel) before (left plot) and after (right plot) the modification of clr$P_j$ values at zero entries of the taxa count table. 'Cor' represents the sample Pearson's correlation coefficient.

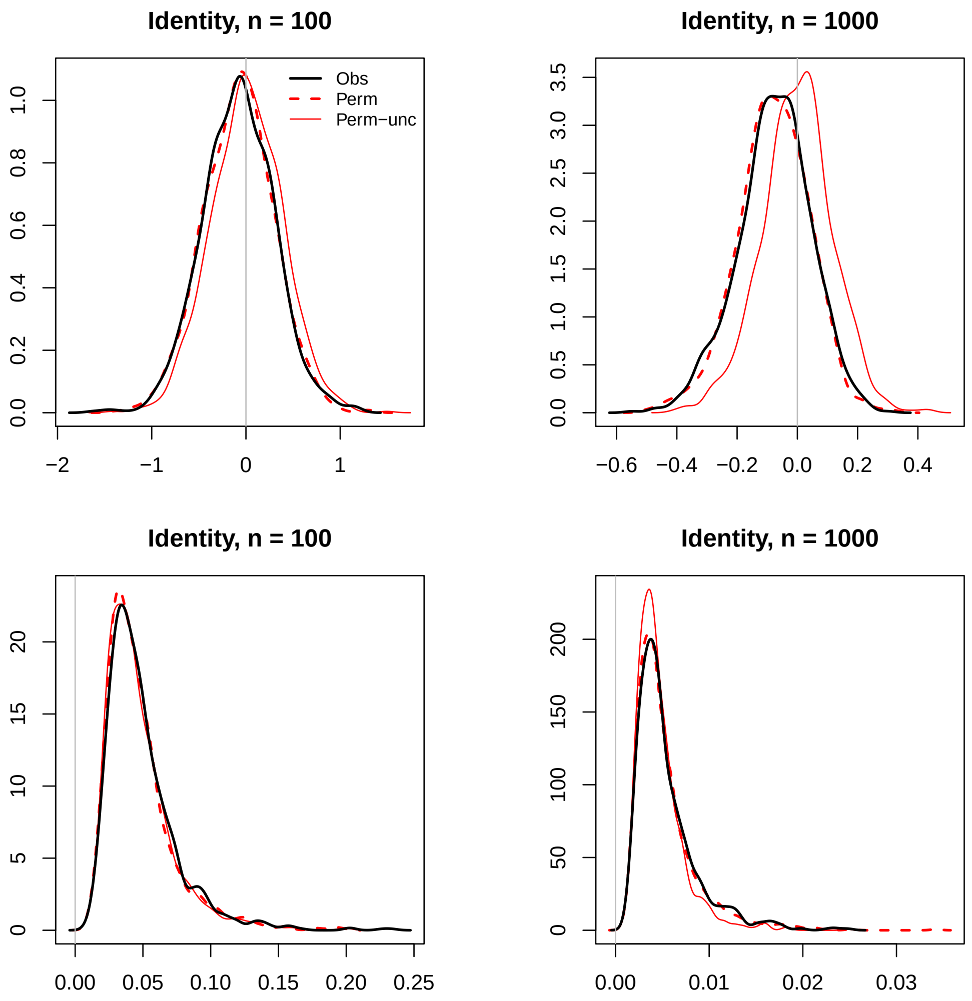


Figure S18: Distributions of sample Pearson's covariances (upper panel) and sample distance covariances (lower panel) across all null taxon pairs, when the covariance structure is Identity. The observed statistics ('Obs', black solid line) are based on one replicate of observed data. The permutation statistics ('Perm', red dashed line) are based on one permutation replicate, for which we also calculated the corresponding permutation statistics without centering $\widetilde{\text{clr}P_j}$ ('Perm-unc', red solid line). The gray vertical line represents the zero reference line.

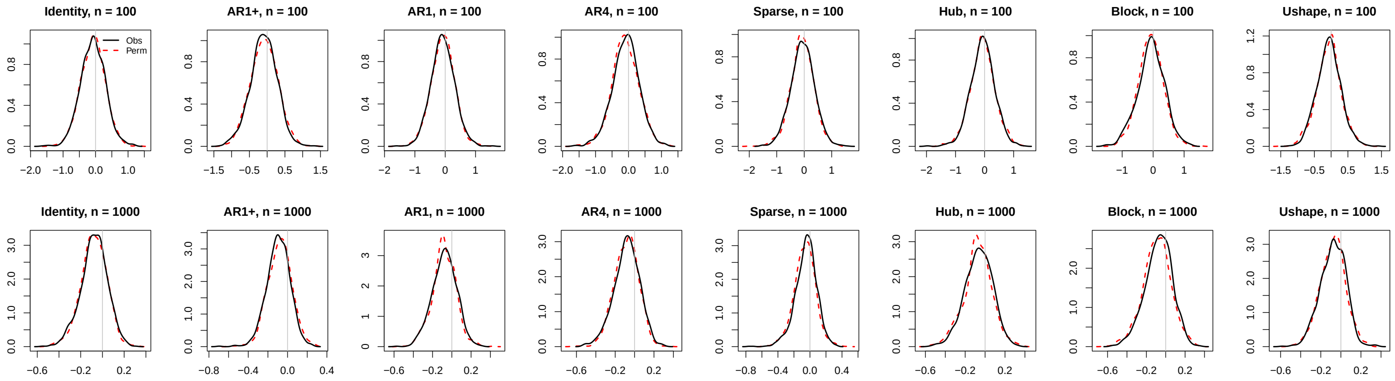


Figure S19: Distributions of samples Pearson's covariances for data filtered with the stringent criterion. Refer to the caption of Figure S18 for additional information.

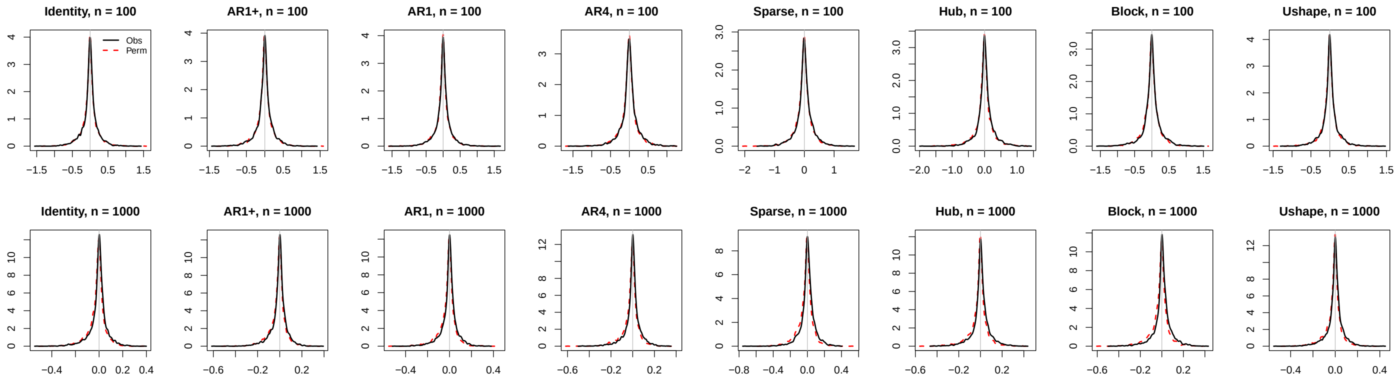


Figure S20: Distributions of sample Pearson's covariances for data filtered with the lenient criterion. Refer to the caption of Figure S18 for additional information.

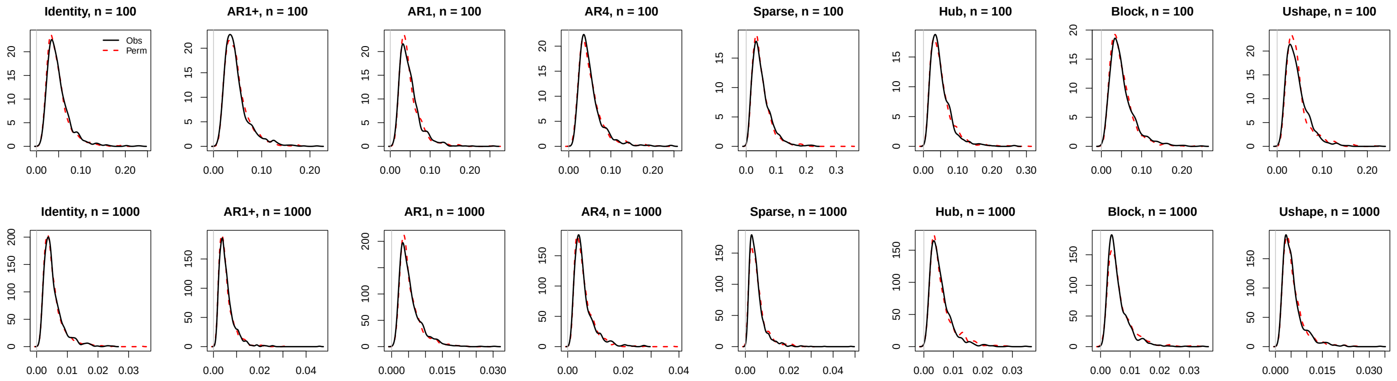


Figure S21: Distributions of sample distance covariances for data filtered with the stringent criterion. Refer to the caption of Figure S18 for additional information.

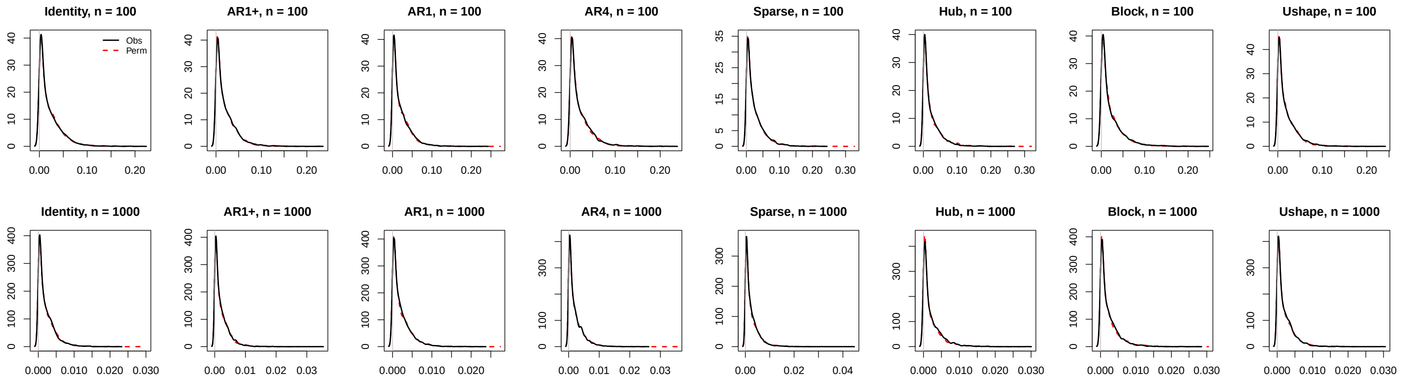


Figure S22: Distributions of sample distance covariances for data filtered with the lenient criterion. Refer to the caption of Figure S18 for additional information.
